# Intracellular Tau Fragment Droplets Serve as Seeds for Tau Fibrils

**DOI:** 10.1101/2023.09.10.557018

**Authors:** Yoshiyuki Soeda, Hideaki Yoshimura, Hiroko Bannai, Riki Koike, Isshin Shiiba, Akihiko Takashima

## Abstract

Intracellular tau aggregation requires a local protein concentration increase, referred to as "droplets". However, the cellular mechanism for droplet formation is poorly understood. Here, we expressed OptoTau, a P301L mutant tau fused with CRY2olig, a light-sensitive protein that can form homo-oligomers. Under blue light exposure, OptoTau increased tau phosphorylation and was sequestered in aggresomes. Suppressing aggresome formation by nocodazole formed tau granular clusters in the cytoplasm. The granular clusters disappeared by discontinuing blue light exposure or 1,6-hexanediol treatment suggesting that intracellular tau droplet formation requires microtubule collapse. Expressing OptoTau-ΔN, a species of N-terminal cleaved tau observed in the Alzheimer’s disease brain, formed 1,6-hexanediol and detergent-resistant tau clusters in the cytoplasm with blue light stimulation. This intracellular stable tau clusters acted as a seed for tau fibrils in vitro. These results suggest that tau droplet formation and N-terminal cleavage are necessary for neurofibrillary tangles formation in neurodegenerative diseases.

## Introduction

Tau, a microtubule-binding protein, is an intrinsically disordered protein that is expressed as six isoforms in the adult human brain. Tau modulates microtubule dynamics and the stabilization of the cytoskeleton through direct association with tubulin dimers ^1,2^. In tauopathies, hyper-phosphorylation of tau reduces the affinity of tau for microtubules ^3^, resulting in microtubule collapse ^4^ ^5,6^. Hyperphosphorylated tau molecules bind together and form fibrillar tau aggregates with β-sheet structure in cells of the central nervous system^7-10^. Recently, cryo-electron microscopy (cryo-EM) studies revealed the atomic structures of tau fibril cores from patients with tauopathies such as Alzheimer’s disease (AD)^11^, Pick’s disease^12^, chronic traumatic encephalopathy^13^, corticobasal degeneration^14,15^, progressive supranuclear palsy^16^, argyrophilic grain disease^16^, and primary age-related tauopathies^16^. These studies suggest that the tau core structure consists of different amino acid sequences of tau in each disease, although this variability is limited in the region from the microtubule-binding domain to the C-terminal domain of tau. However, the mechanism of formation of ordered aggregate tau fibrils from the highly disordered protein tau is unclear.

In previous studies, the oligomerization of tau preceding the formation of tau fibrils has been described as an early event in the pathogenesis of tauopathies ^17,18^ ^19^ and to be involved in toxicity ^20-24^. Our in vitro studies using recombinant tau have shown that tau fibrils consist of granular tau oligomers^25^, which have been observed in the frontal cortex of Braak stage 1 patients^26^. As a mechanism for tau oligomer formation, recent reports have shown that recombinant tau first undergoes liquid-liquid phase separation (LLPS) by exposure to a crowding agent and polyanions, forming condensates referred to as tau droplets ^27-31^. Droplet tau transitions into aggregates with β-sheet structure through long incubation ^30,31^, and the process of aggregation has been clarified in vitro. However, the intracellular mechanism of tau aggregation, including tau droplet formation and transition from tau droplet to tau oligomer formation, is not fully characterized.

Arabidopsis cryptochrome 2 (CRY2) is a photoreceptor-soluble protein, that modulates circadian rhythm in plants, leading to the regulation of development and growth. The CRY2 protein, which can undergo homo-oligomerization under blue light, is a powerful optogenetic tool for inducing the intracellular condensation of target proteins by blue light in mammalian cells ^32,33^ ^34^. CRY2olig is a CRY2 mutant (E490G) with increased oligomerization capacity compared to CRY2 ^32,35^. CRY2olig oligomers formed using blue light can be dissociated by light deprivation ^32^. Amyloid proteins, including α-synuclein ^36^ and TAR DNA-binding protein of 43 kDa (TDP-43) ^37,38^, when fused to CRY2olig, form oligomers and become supersaturated upon exposure to blue light in vivo and in vitro. Thus, CRY2olig can be used to analyze the cellular processes involved in tau protein condensation through aggregation with β-sheet structure.

In the present study, we used CRY2olig fused to tau to investigate the conditions under which tau assembles to form intracellular droplets within the cell and, then, the conditions under which tau droplets transition to tau oligomers.

## Results

### Full-length-P301L with CRY2olig (OptoTau) clusters in microtubules and is transported to aggresomes

We used the frontotemporal dementia with parkinsonism (FTDP)-17 mutation P301L mutant tau fused with CRY2olig (OptoTau; Fig. S1) in all experiments, because P301L has been reported to be a tau aggregation-prone mutation ^21,39^. Western blot analyses of OptoTau expressed in the mouse neuroblastoma cell line, Neuro2a, indicated that OptoTau (150 kDa) showed a mobility shift when exposed to blue light for 24 hrs (Fig. 1A), suggesting an increase in OptoTau phosphorylation by blue light exposure. This was confirmed by the elevation of tau phosphorylation at the AT8 site (Fig. 1B, left panel) and S422 (Fig. 1B, right panel), which are involved in tau aggregation ^40-42^, S262 (Fig. S2A), which is involved in microtubule binding ^43^, and T217 (Fig. S2B), which is a biomarker of AD ^44,45^. Expressing tau without CRY2olig did not show blue light-dependent tau phosphorylation (Fig. S3), which suggests that enforced clustering of tau in the cells increased tau phosphorylation. Further, quantification of reactive oxygen species (ROS) using CellROX Green Reagent showed no significant increase in ROS levels after 24 hrs of light exposure (Fig. S10), indicating that, under the conditions used in our experiments, there was no phototoxicity in terms of ROS generation. In non-reducing western blot analyses, we observed a significant (9.7-fold) increase in the tau band above 250 kDa with blue light exposure compared with that observed after no blue light exposure (Fig. 1C). These results suggest that the clustering of OptoTau leads to tau hyper-phosphorylation and oligomerization.

**Figure 1:**
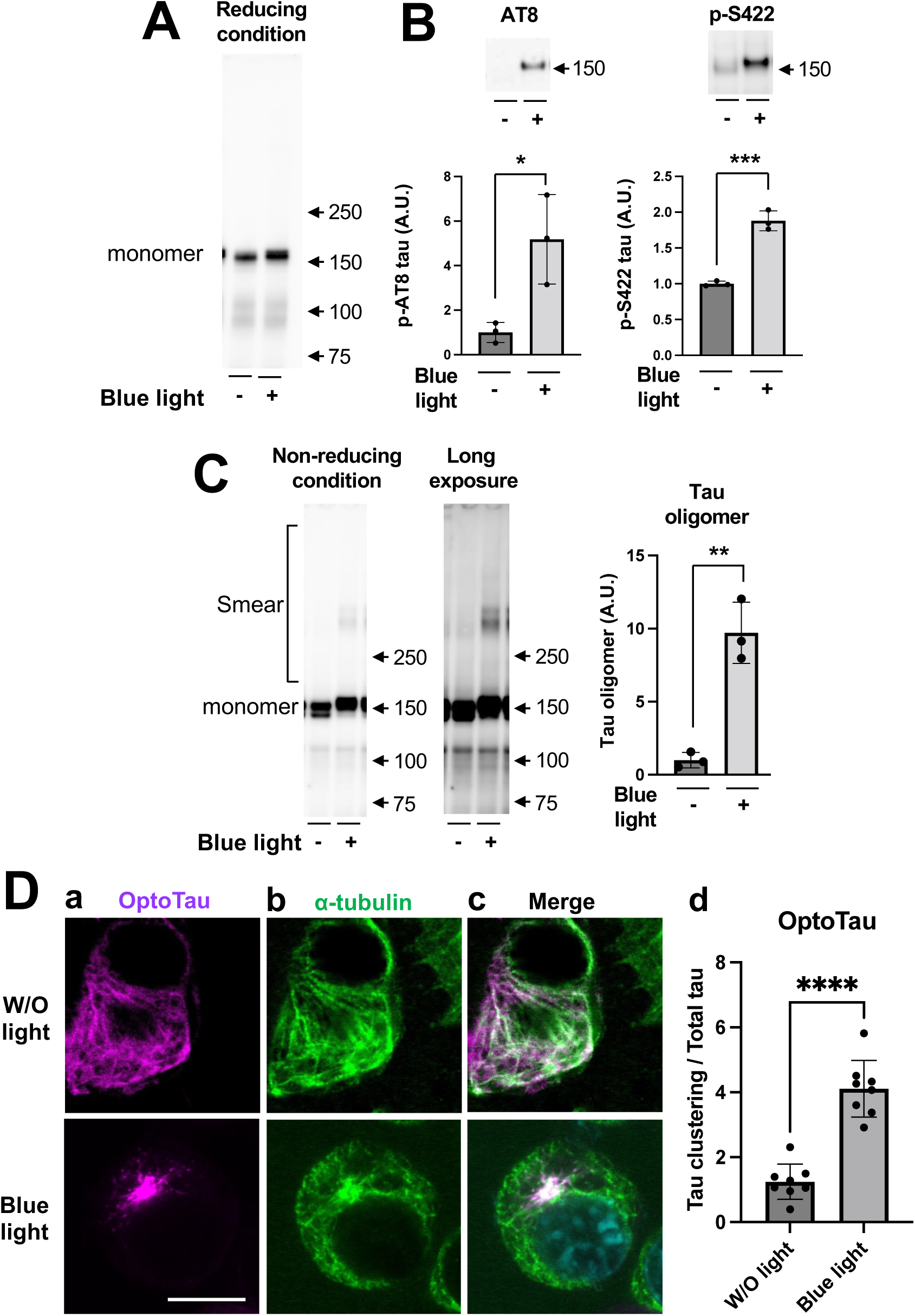
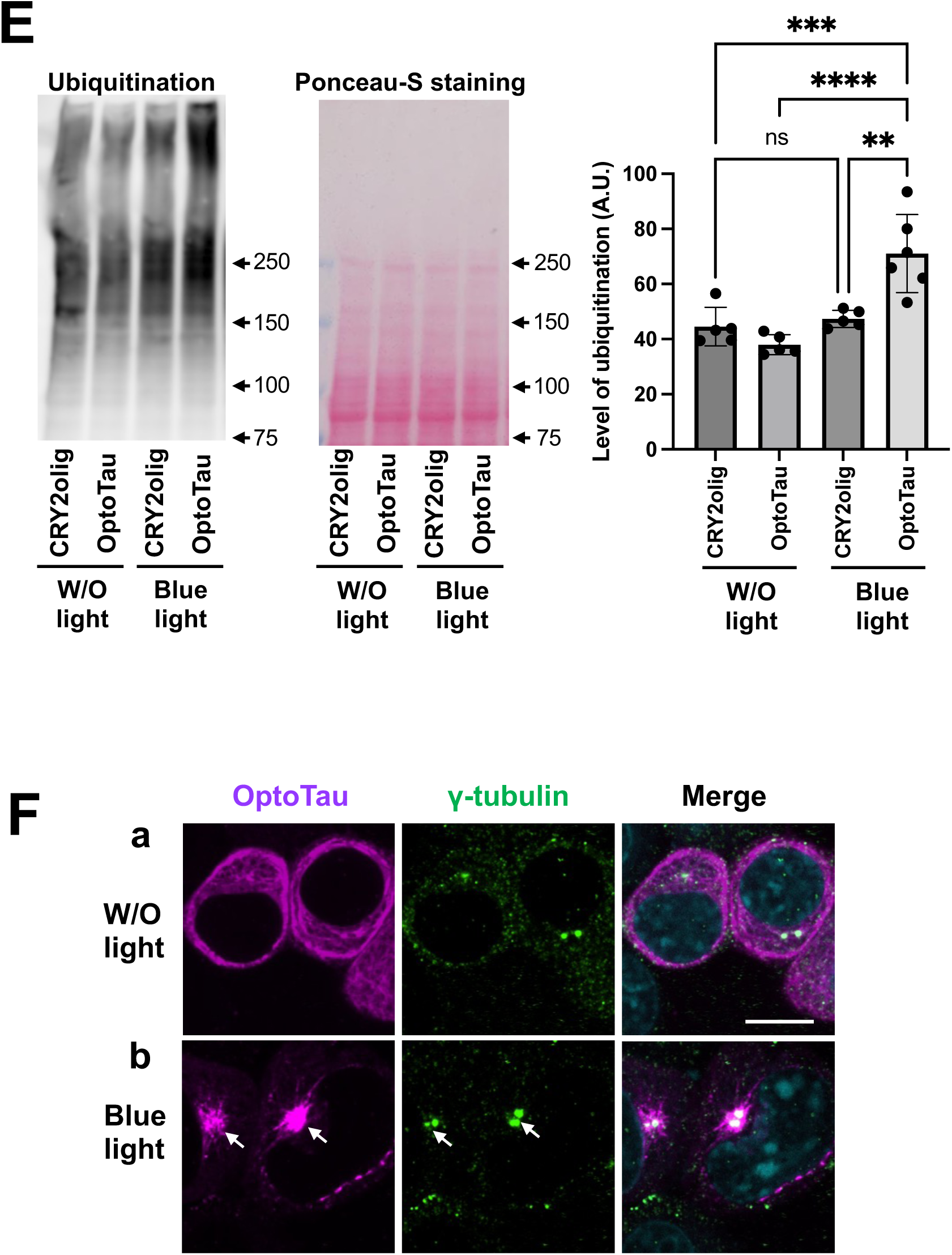
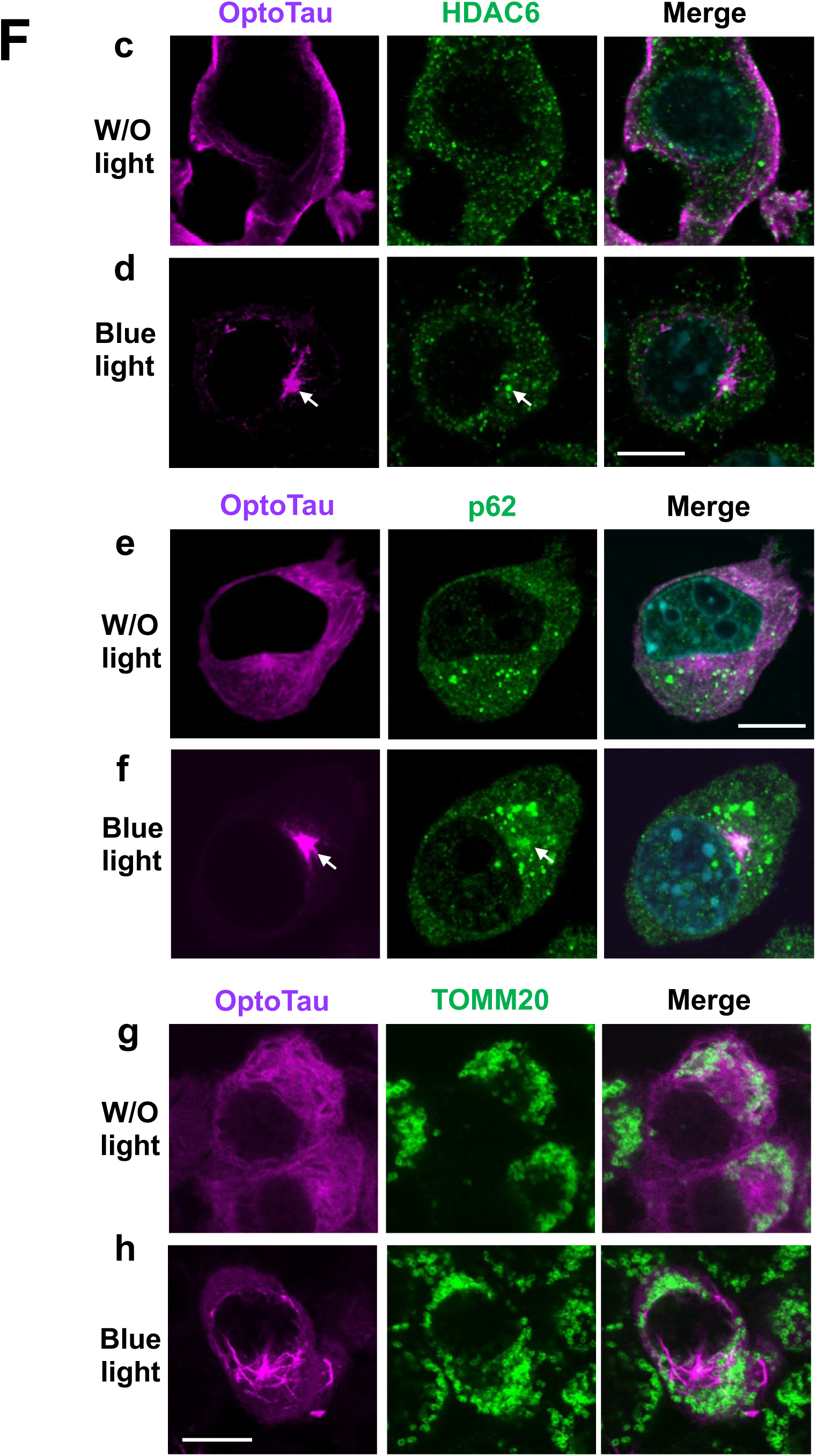
OptoTau is clustered on microtubules and transported to aggresomes. (A,B) Under reducing conditions, tau protein was detected by pan-tau JM antibody (A), and phospho-tau was detected by AT8 antibody, which recognizes tau phosphorylation at S202 and T205 (B, left panel) and by p-S422-tau antibody (B, right panel) in Neuro2a cells expressing OptoTau with or without exposure to blue light for 24 hrs. Levels of phospho-tau were normalized to corresponding total tau (B, lower panel). (C) Under non-reducing conditions, the level of tau protein was determined by SDS-PAGE western blotting using JM antibody. To detect the tau smear band on the membranes clearly, a long period of exposure was employed (C, middle panel). The tau smear bands were quantified (C, right panel). (D) Colocalization of OptoTau in microtubules. The cells were stained with SNAP-Cell TMR-Star to detect OptoTau (magenta) (a), and immuno-stained with α-tubulin antibody (green) (b) at 24 hrs after blue light exposure. Images of OptoTau and α-tubulin are merged (c). (d) OptoTau intensity in the images was quantified using ImageJ. During blue light exposure, regions of interest (ROIs) were defined as regions where OptoTau clustering was observed, and their intensity was measured. In the W/O light exposure, ROIs were set around the nucleus, and their area was the same as under blue light conditions. No difference in ROIs area was observed between blue light exposure and W/O light exposure. (Fig. S11). Total OptoTau was quantified by measuring the intensity of OptoTau across the entire cell. The OptoTau clustering intensity was normalized by the total OptoTau intensity. (E) Ubiquitinated proteins were detected (E, left panel) and quantified (E, right panel) in Neuro2a cells expressing OptoTau and CRY2olig control with or without blue light exposure for 24 hrs. Total proteins were stained by Ponceau-S (E, middle panel). (F) Colocalization of OptoTau in aggresomes. After exposure with blue light or W/O light for 20-24 hrs, the cells were stained with SNAP-Cell TMR-Star (F, left panel, white arrow), and immuno-stained with γ-tubulin (a,b), histone deacetylase 6 (HDAC6) (c,d) and p62 (e,f) TOMM20 (g,h) (green) (F, middle panel, white arrow). Images are merged (F, right panel). All error bars indicate mean ± SD. *p < 0.05, ** < 0.01, *** < 0.001 and **** < 0.0001 by Student’s t-test (B,C,D) and one-way analysis of variance and Tukey’s multiple comparisons test (E). p-, phosphorylated; W/O, without. The white bar is 10 μm.

To investigate the localization of CRY2-FL-tau clusters in the cells, we performed co-immunofluorescence staining for OptoTau and α-tubulin. Using confocal microscopy and reflected light microscopy, OptoTau was observed to colocalize with α-tubulin and concentrate near the nucleus in blue light-exposed cells, while, in the absence of light exposure, OptoTau distributed uniformly throughout the cells, along with and without α-tubulin (Figs. 1D and S4A). The blue light-induced OptoTau clustering structure may correspond to aggresomes, which are generated near or surrounding the cell’s centrosomes^46^. The protein becomes polyubiquitinated, allowing for retrograde transportation facilitated by the binding of histone deacetylase 6 (HDAC6) ^46,47^. Aggregates of p62-bound ubiquitinated proteins interact with HDAC6 and are transported to the microtubule-organizing center, where they form aggresomes^48^ ^49^. The aggresomes are surrounded by mitochondria^50^. In blue light-exposed cells, overexpressed OptoTau induced increased levels of ubiquitinated protein, compared with the CRY2olig control (Fig. 1E). Furthermore, γ-tubulin, HDAC6, and p62, but not the mitochondria marker TOMM20 colocalized with OptoTau (Figs. 1F, S9C, S9D, S9E and S9F). TOMM20 appears to surround the OptoTau (Fig. 1F). These results suggest that clustering OptoTau migrates along microtubules to aggresomes and may be degraded. Thus, we investigated the effect of inhibiting aggresome formation through the destruction of microtubules.

### Blue light exposure induces OptoTau droplet formation even with inhibition of microtubule assembly

To inhibit microtubule polymerization, Neuro2a cells expressing OptoTau were treated with nocodazole, a reversible microtubule polymerization inhibitor ^51^, during blue light exposure (Fig. 2A). As shown in Figures 2B-D, blue light exposure to nocodazole-treated cells induced an increased tau phosphorylation (Fig. 2C) and oligomer formation (Fig. 2D). Thus, the evoked stimulation of tau phosphorylation and tau oligomerization by blue light exposure was maintained even with nocodazole treatment. Fluorescence microscopy revealed that the granular OptoTau clusters were localized not only in the perinuclear area, but also throughout the cell (Figs. 2E, S4B, and S5). Further, the granular OptoTau clusters and γ-tubulin were not observed near or surrounding the cell’s centrosomes (Figs. 2F and S9H), suggesting that microtubule destruction led the clustering tau to form membrane free structures in the cytoplasm, but not aggresomes.

**Figure 2:**
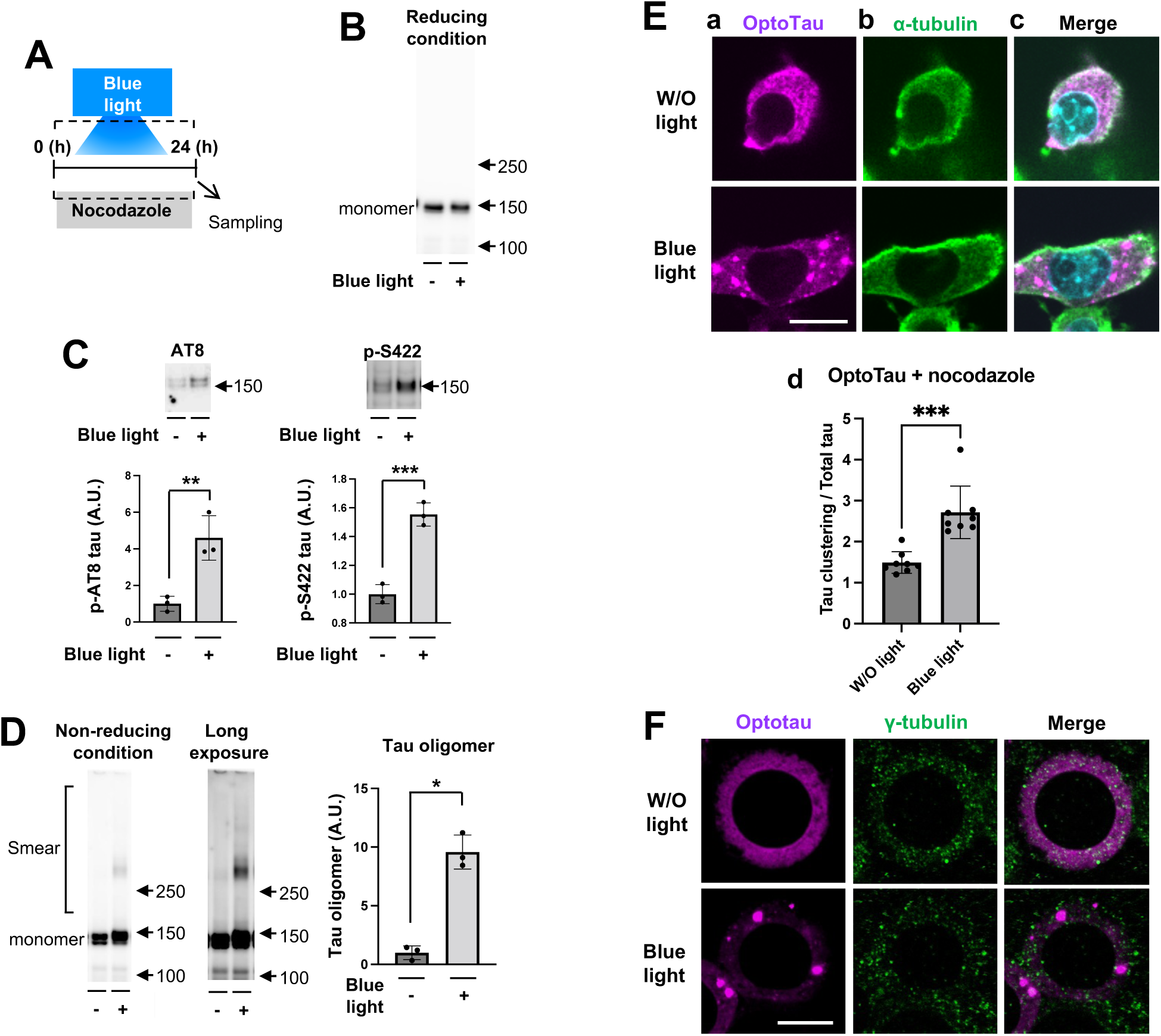
OptoTau is concentrated in the cytosol by the inhibition of microtubule assembly. (A) Scheme of nocodazole exposure. (B,C) After microtubules were depolymerized by nocodazole, tau protein was detected by pan-tau JM antibody (B) and phospho-tau was detected by AT8 antibody (C, left panel) and by p-S422-tau antibody (C, right panel) in Neuro2a cells expressing OptoTau with or without blue light exposure for 24 hrs. Levels of phospho-tau were normalized by corresponding total tau (C, lower panel). (D) Under non-reducing condition, the level of tau protein was determined by SDS-PAGE western blotting using JM antibody. (E,F) The clustering of OptoTau under exposures of nocodazole and blue light for 24 hrs was not colocalized with aggresomes. The cells were stained with SNAP-Cell TMR-Star to detect OptoTau (magenta) (E-a and F, left panel), and immuno-stained with α-tubulin antibody (E-b) and γ-tubulin (F, middle panel)(green). Images were merged (E-c and F, right panel). (E-d) OptoTau intensity in the images was quantified using ImageJ. The OptoTau clustering intensity was normalized by the total OptoTau intensity. All error bars indicate mean ± SD. *p < 0.05, ** < 0.01 and *** < 0.001 by Student’s t-test. p-, phosphorylated. W/O, without. The white bar is 10 μm.

To further characterize the granular tau clustering, we removed the cells from blue light exposure and counted the number of the granular tau clusters. The absence of blue light significantly decreased the number of granular OptoTau clusters (Fig. 3A), which indicates that the granular OptoTau clustering by nocodazole and blue light exposure is reversible, consistent with the results of previous studies ^27-31^.

**Figure 3:**
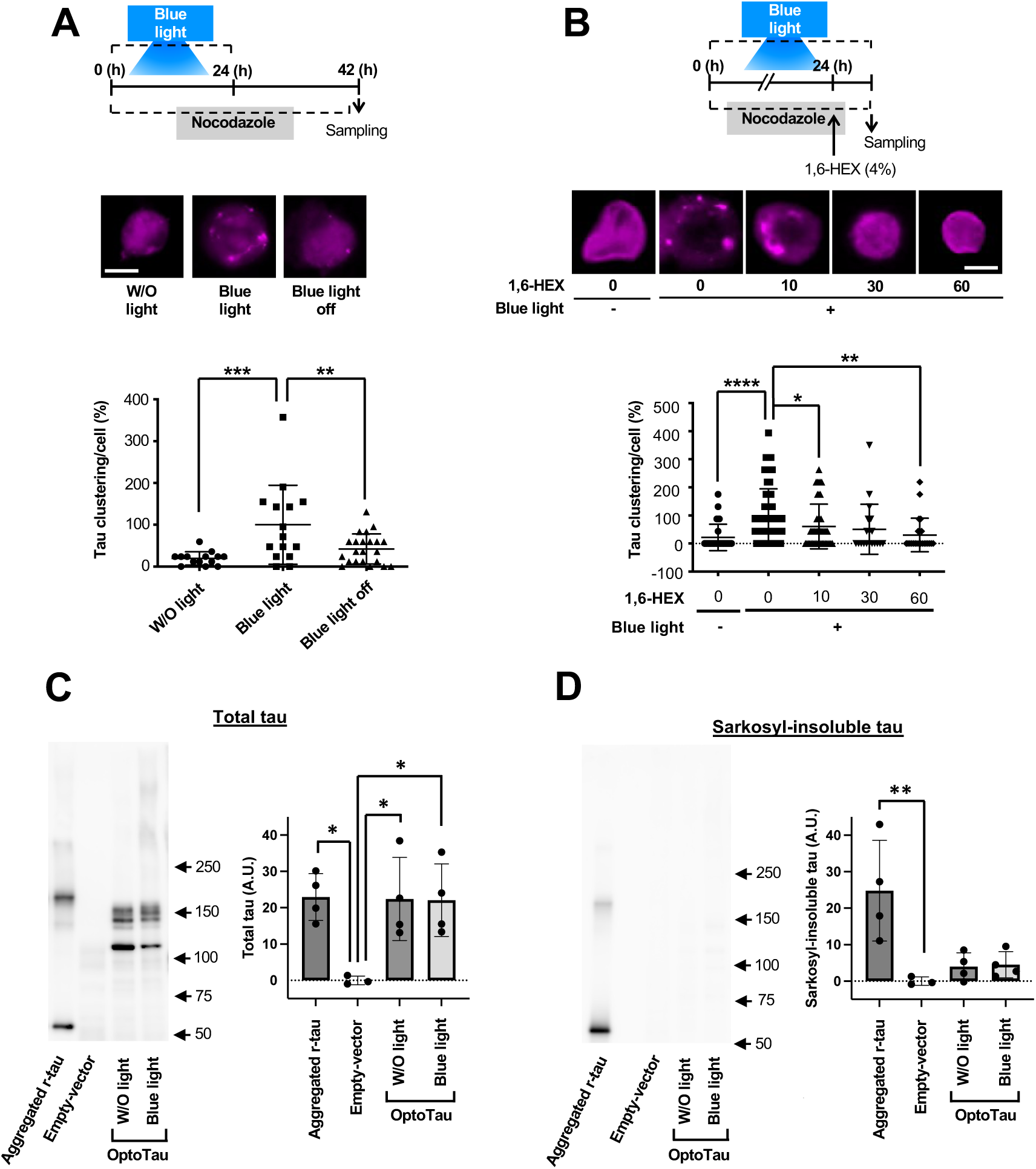
The reversible tau clusters in cells expressing OptoTau. (A) Blue light was turned off after the cells expressing OptoTau were exposed for 24 hrs in the presence of nocodazole. The cells were stained with SNAP-Cell TMR-Star at 18 hrs after the blue light was turned off. (B) 1,6-hexanediol (1,6-HEX) was added to the cells expressing OptoTau under exposures of nocodazole and blue light for 24 hrs. (C-D) Clustering OptoTau under exposures of nocodazole and blue light for 23 hrs was solubilized with anionic surfactant sarkosyl. After homogenates from cells expressing OptoTau and an empty vector, and aggregated recombinant tau were incubated with 1% sarkosyl for 1 hr at 37C and centrifuged. The levels of tau protein in the total fraction (C) and sarkosyl-insoluble fraction (D) were determined by SDS-PAGE western blotting using JM antibody (left panel). The tau bands were quantified (right panel). All error bars indicate mean ± SD. *p < 0.05, ** < 0.01, *** < 0.001 and **** < 0.0001 by one-way analysis of variance and Dunnett’s multiple comparisons test. The white bar is 10 μm. W/O, without; r-, recombinant.

Protein droplets and membrane-less organelles can be dissolved in vitro and in cellular studies using 1,6-hexanediol, but stable protein aggregations including the tau fibril, remain intact in the presence of 1,6-hexanediol ^52-55^ ^56^. Blue light-induced granular OptoTau clustering completely disappeared by treatment with 1,6-hexanediol for 60 minutes (Fig. 3B, Supplemental video 1). However, the accumulation of aggregated tau exhibits resistance to the anionic surfactant sarkosyl ^57^. While aggregated recombinant tau as a positive control was insoluble with sarkosyl, the sarkosyl-insoluble tau was rarely detected in the OptoTau-expressing cell lysates containing granular tau clusters (Fig. 3D). No difference in the amount of total tau was detected between the samples (Fig. 3C). Taken together, OptoTau forms a membrane-less structure of a droplet-like nature in the cell when exposed to blue light under the condition of nocodazole treatment.

### OptoTau carrying an N-terminal deletion (OptoTau-ΔN) is aggregated by blue light in the cytosol of cultured cells

Microtubule disruption could form from the OptoTau clustered to an intracellular droplet, but this condition could not form a stable tau aggregation. An additional condition is required for tau aggregation formation from the tau droplets. Cryo-EM studies revealed that, in tauopathies, the core of tau fibrils is composed of a common sequence, including a microtubule-binding repeat domain and part of the C-terminal domain ^11^ ^12^ ^13^ ^14,15^ ^16^. Deletion of the N-terminal projection domain increases with aging in tau transgenic mice, and accelerates heparin-induced tau aggregation in vitro ^58^. These findings suggested that we investigate the effects of N-terminal deletion in the tau aggregation process. We generated OptoTau-ΔN using CRY2olig joined with P301L carrying an N-terminal deletion (Fig. 4A). OptoTau-ΔN expression showed a tau band mobility shift from that of OptoTau and an increase of tau phosphorylation in response to blue light exposure (Fig. 4B). Further, an 8.3-fold increase in tau oligomerization was observed in cells exposed to blue light (Fig. 4C), indicating that the N-terminal deletion does not affect the tau clustering process. Under fluorescence microscopy observation, OptoTau staining showed the assembly of tau clusters and not granular tau (Figs. 4D and S4C). Sarkosyl-insoluble tau was significantly increased in cells expressing OptoTau-ΔN when exposed to blue light compared with cells expressing OptoTau (Fig. 4E). Further, the blue light-induced OptoTau-ΔN clusters were colocalized with Thioflavin S-positive staining (Figs. 4F and S9K). These results indicate that OptoTau-ΔN is capable of facilitating tau aggregation containing a β-sheet structure. After turning off the blue light (Fig. 4G) or exposure to 1,6-hexanediol (Fig. 4H, Supplemental video 2), OptoTau-ΔN clusters remained stable, unlike the tau droplet. These results suggest that N-terminal deletion enhances tau-tau interaction, leading to the transition of tau droplets to stable tau aggregates.

**Figure 4:**
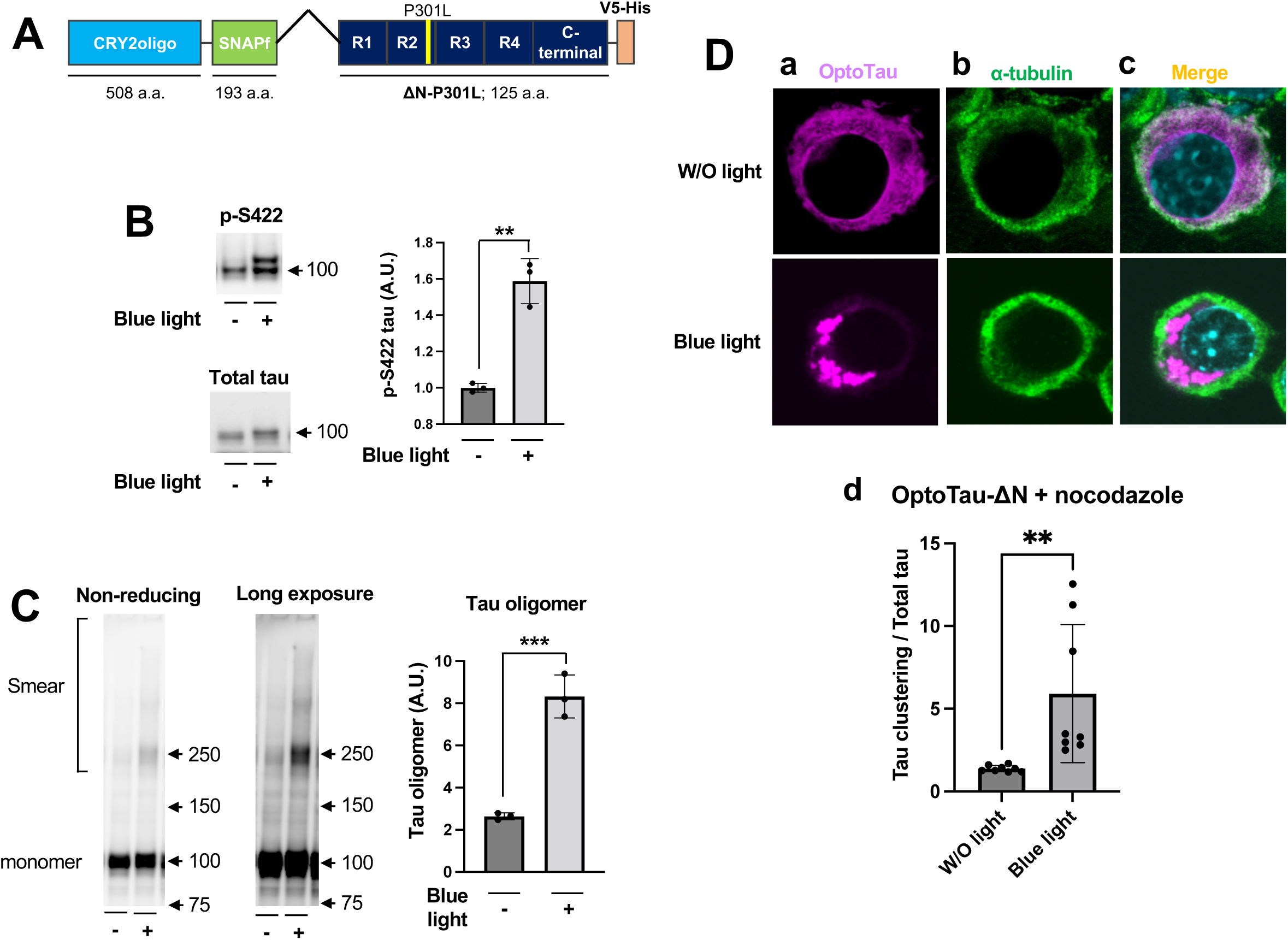
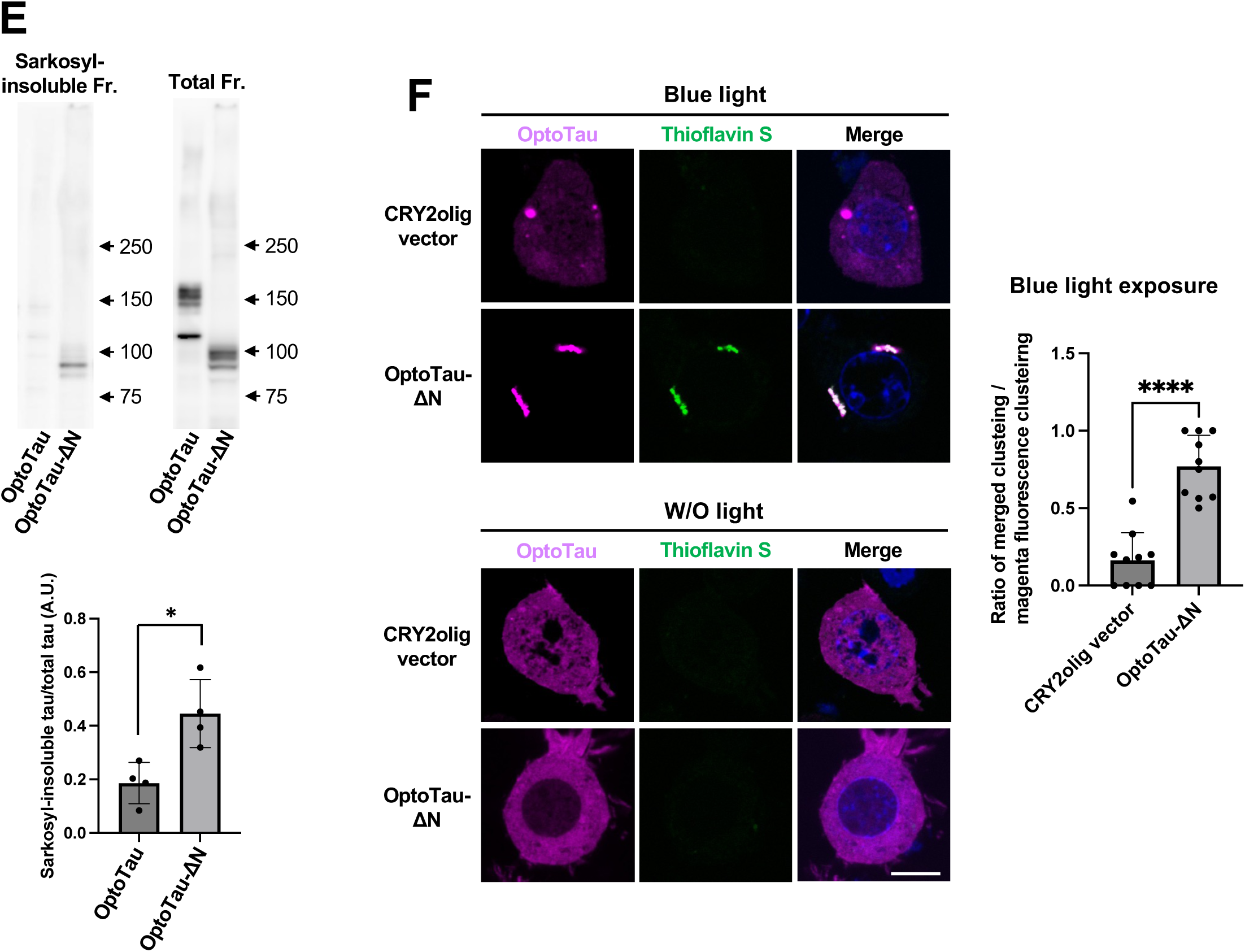
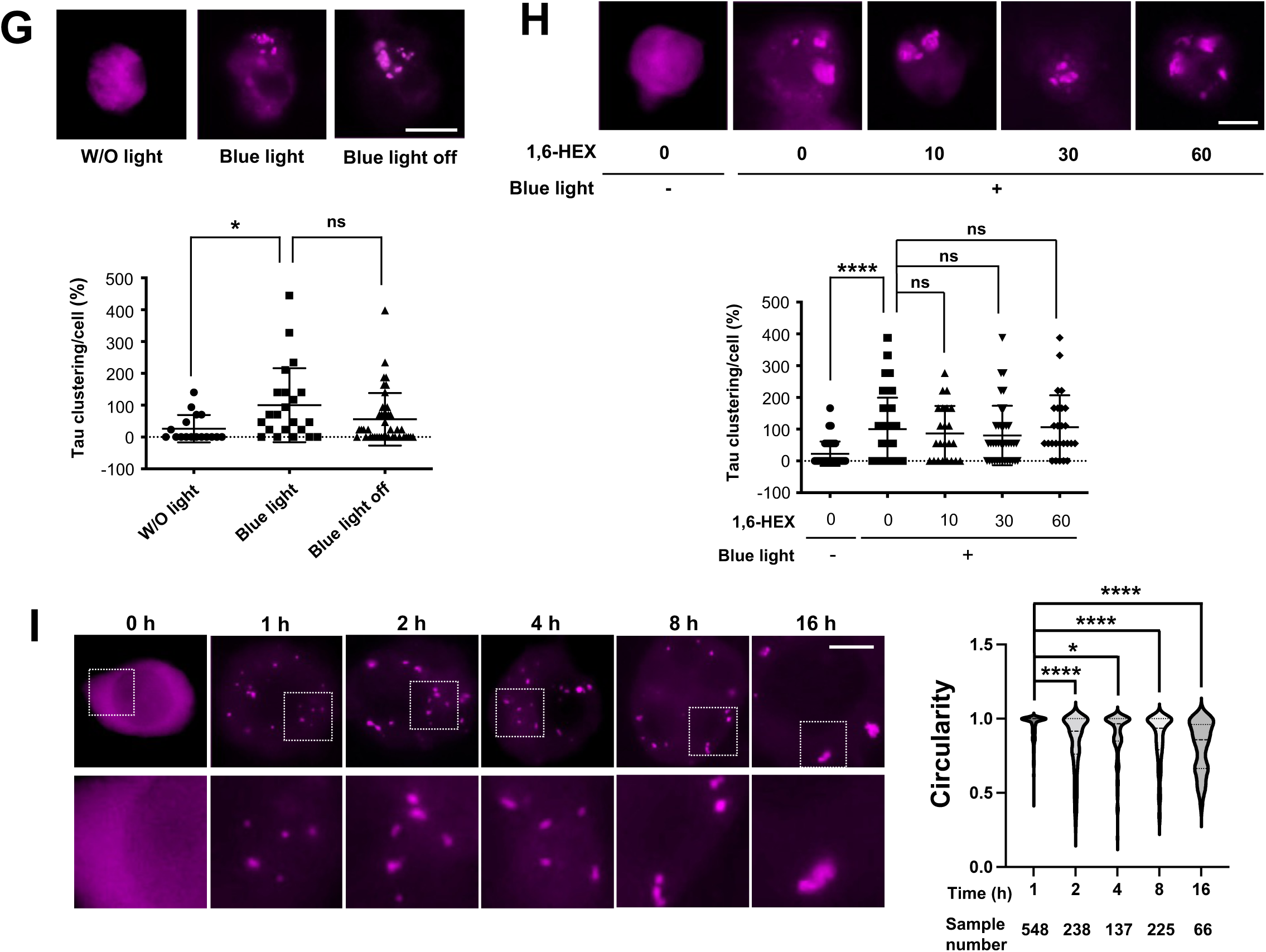
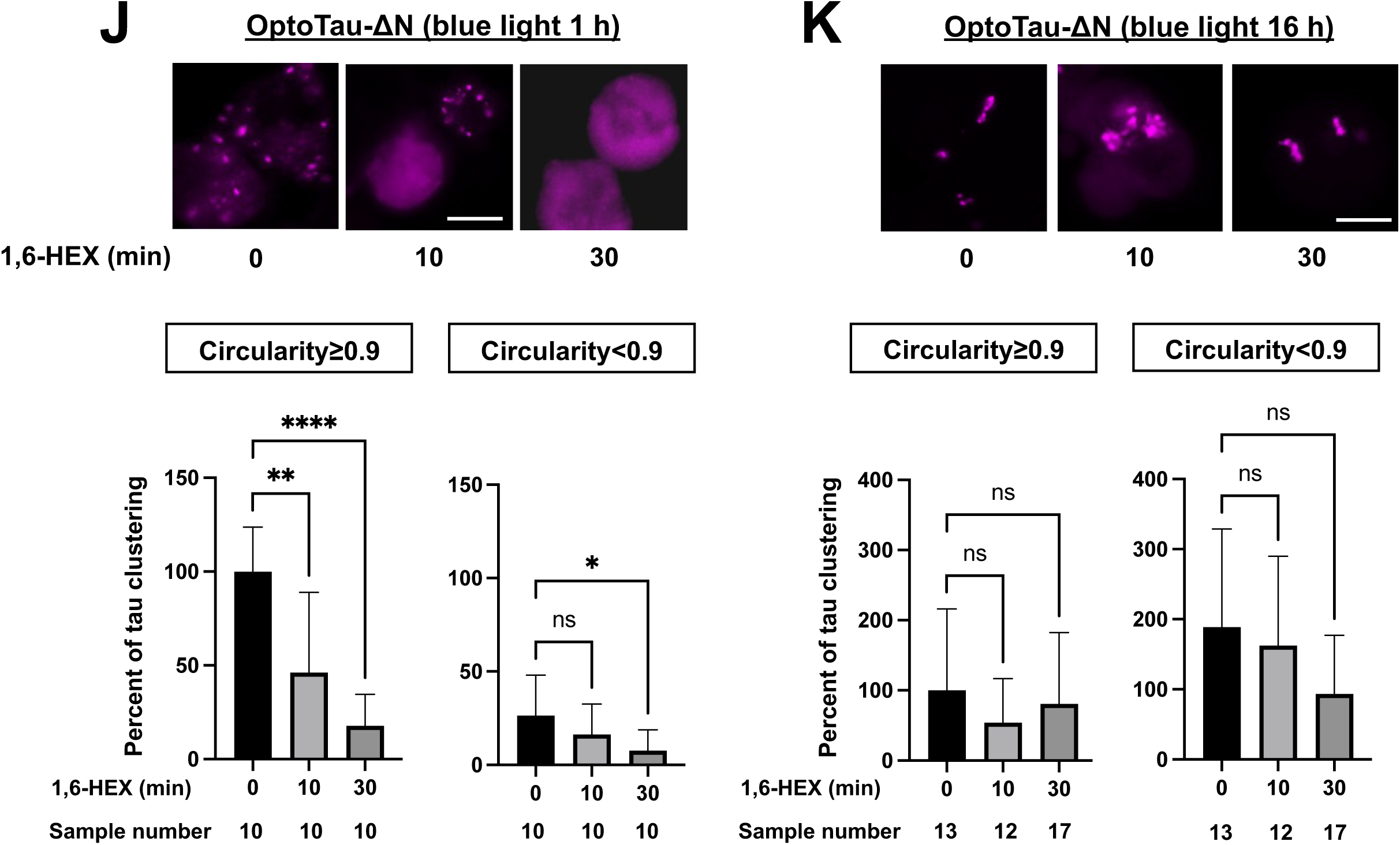
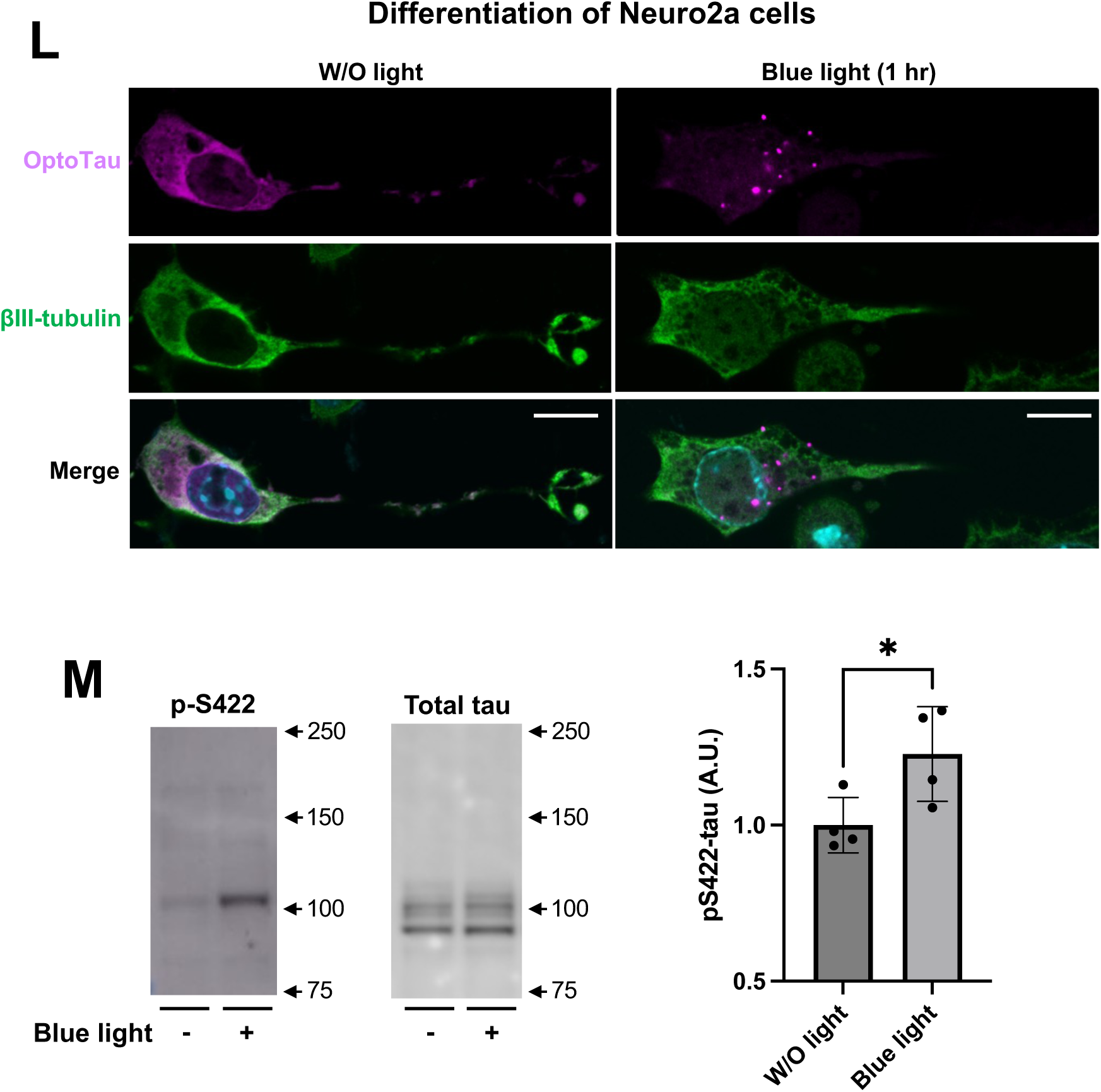
The irreversible tau clusters in cells expressing OptoTau-ΔN. (A) Structure of OptoTau-ΔN. (B) OptoTau-ΔN-expressing Neuro2a cells were exposed to nocodazole and blue light for 24 hrs, then phospho-tau was detected by p-S422-tau antibody (left and upper panel) and tau protein was detected by pan-tau JM antibody (left and lower panel) using reducing condition SDS-PAGE western blotting. The level of phospho-tau was normalized to the corresponding total tau (right panel). (C) Under non-reducing condition, the level of tau protein was determined by SDS-PAGE western blotting using JM antibody. (D) The cells were stained with SNAP-Cell TMR-Star to detect OptoTau-ΔN (magenta) (a), and immuno-stained with α-tubulin antibody (green) (b) at 24 hrs after exposure to nocodazole and blue light. Images are merged (c). (d) OptoTau-ΔN intensity in the images was quantified using ImageJ. The OptoTau-ΔN clustering intensity was normalized by the total OptoTau-ΔN intensity. (E) OptoTau-ΔN was more insoluble with sarkosyl than OptoTau. Cells were collected and homogenized after exposure to nocodazole and blue light for 23 hrs. The cell homogenates were incubated with 1% sarkosyl for 1 hr at 37C. The levels of tau protein in the sarkosyl-insoluble fractions (left panel) and the cell homogenates (total fr., middle panel) were determined by SDS-PAGE western blotting using JM antibody. The tau bands were quantified, and levels of sarkosyl-insoluble tau were normalized by corresponding total tau (right panel). (F) The cells were transfected with OptoTau-ΔN and CRY2olig vector. After exposure to blue light for 18 hrs, the cells were stained with SNAP-Cell TMR-Star (magenta), and thioflavin S (green). Under exposure of blue light, the number of clusters bearing merged signals was normalized to the corresponding clusters bearing magenta signals. (G,H) The clustering of OptoTau-ΔN under exposure to nocodazole and blue light grows into an irreversible aggregated structure. After the exposure for 24 h, the light was turned off for 18 hrs (G) or the cells were treated with 1,6-hexanediol (1,6-HEX) (H). OptoTau-ΔN in the cells was stained with SNAP-Cell TMR-Star as magenta fluorescence. (I) The circularity of tau clustering was time-dependently reduced by blue light exposure. The cells expressing OptoTau-ΔN were exposed to blue light and 10 μM nocodazole at the indicated time, and stained with SNAP-Cell TMR-Star. The lower panels show expanded images in the dotted white square. (J,K) The structure formed by the blue light short time exposure is a droplet formed by LLPS. The cells expressing OptoTau-ΔN were exposed to nocodazole and blue light for 1 (J) or 16 (K) hrs, followed by incubation with 1,6-HEX for 0-30 minutes. The OptoTau-ΔN clustering was stained with SNAP-Cell TMR-Star. The distribution of circular structures with a threshold of 0.9 and above (left panels) and non-circular structures with a threshold below 0.9 (right panels) was quantified using ImageJ. OptoTau-ΔN clustering with circularity ≥0.9 under no-treatment and 1,6-HEX is expressed as 100%. (L,M) Neuro2a cells were transfected with OptoTau-ΔN and differentiated by retinoic acid. The cells were exposed to blue light and nocodazole for 1 hr. (L) The cells were stained with SNAP-Cell TMR-Star (magenta) and immunostained with antibody for β-III-tubulin, a differentiation marker (green). (M) Tau phosphorylation at S422 residues was detected with reducing condition SDS-PAGE western blotting. All error bars indicate mean ± SD. *p < 0.05, ** < 0.01, *** < 0.001 and **** < 0.0001 by Student’s t-test (B,E,F,M), Welch’s t-test (C) or one-way analysis of variance and Dunnett’s multiple comparisons test (G-K). p-, phosphorylated; W/O, without; Fr, fraction. The white bar is 10 μm.

Next, we investigated the shape of OptoTau-ΔN cluster formation after blue light exposure. At 1hr after blue light exposure, circular-shaped tau clusters appeared in the cytoplasm. The circularity of these tau clusters was mainly 1.0 (Fig. 4I, J). Longer exposure of blue light resulted in the reduction of the circularity of OptoTau-ΔN clusters (Fig. 4I). OptoTau-ΔN clusters formed by the short blue light exposure were solubilized by 1,6-hexanediol (Fig. 4J), whereas the non-spherical OptoTau clusters formed by the long exposure were resistant to 1,6-hexanediol (Fig. 4K). These results suggest that, over time, OptoTau-ΔN clusters convert from a droplet-like to a solid-like state of tau aggregation in response to blue light exposure since the shape of the liquid phase is generally governed by surface tension and becomes spherical ^59-61^, while the shape of the solid phase, such as aggregated, is non-spherical ^62,63^. In differentiated Nuero2a cells expressing OptoTau-ΔN, exposure to blue light and nocodazole for 1 hr induced formation of granular tau clustering (Fig. 4L). Further, tau phosphorylation at S422 was increased 1.23-foldwhen exposed to blue light, compared with the absence of light (Fig. 4M).

To characterize the solid-like state of tau clusters, we examined the cell lysate by sucrose density gradient centrifugation. This method can separate tau aggregated samples from mouse ^64^ and human brains ^26^, as well as recombinant tau ^25^ into fractions based on their aggregate size and composition: Fraction1, no apparent aggregates; Fraction 2, small tau granules; Fraction 3, granular tau oligomers; and Fractions 4-6, short and long tau fibrils. Cell lysate expressing OptoTau-ΔN after treatment with nocodazole and blue light exposure was fractionated by sucrose density gradient centrifugation. Tau levels in Fractions 2 (Fig. 5B) and 3 (Fig. 5C) were significantly increased in cells expressing OptoTau-ΔN by blue light exposure, compared with cells not exposed to blue light, but not in Fraction 1 (Fig. 5A), 4 (Fig. 5D), and 5 (Fig. 5E). These results demonstrate that aggregates from OptoTau-ΔN under conditions of microtubule destruction consist of small tau granules and granular tau oligomer, which is detergent insoluble^25^.

**Figure 5:**
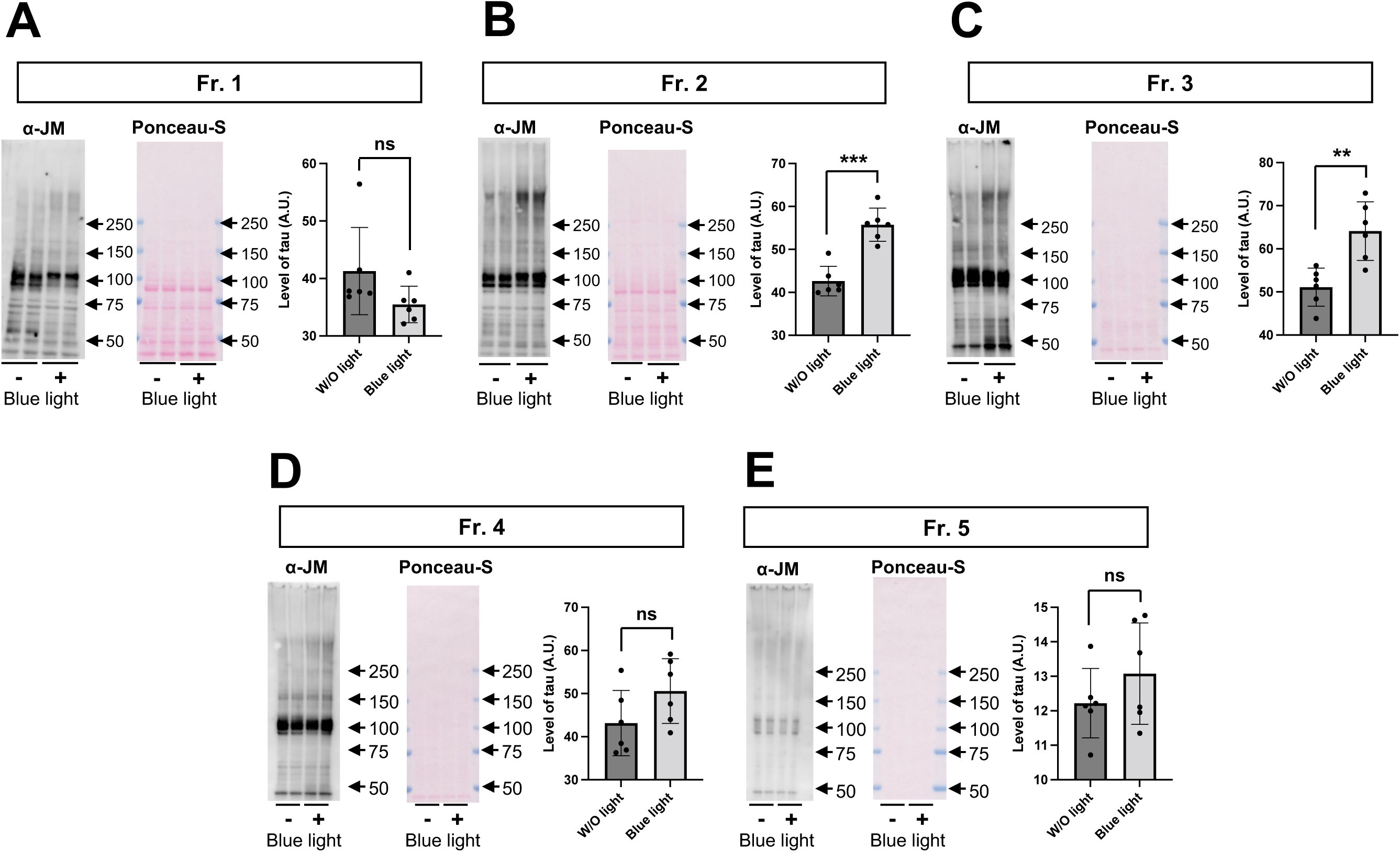
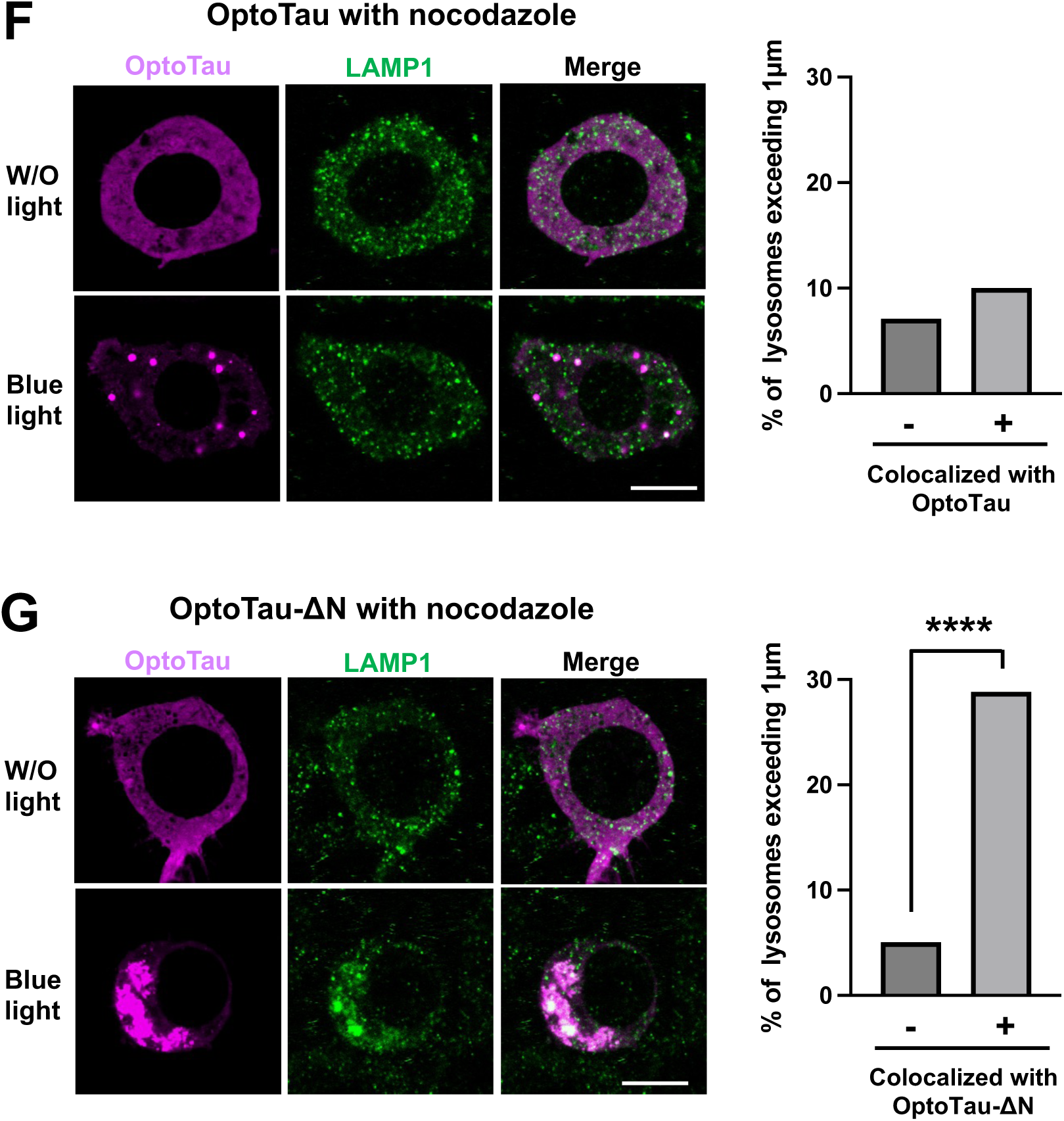
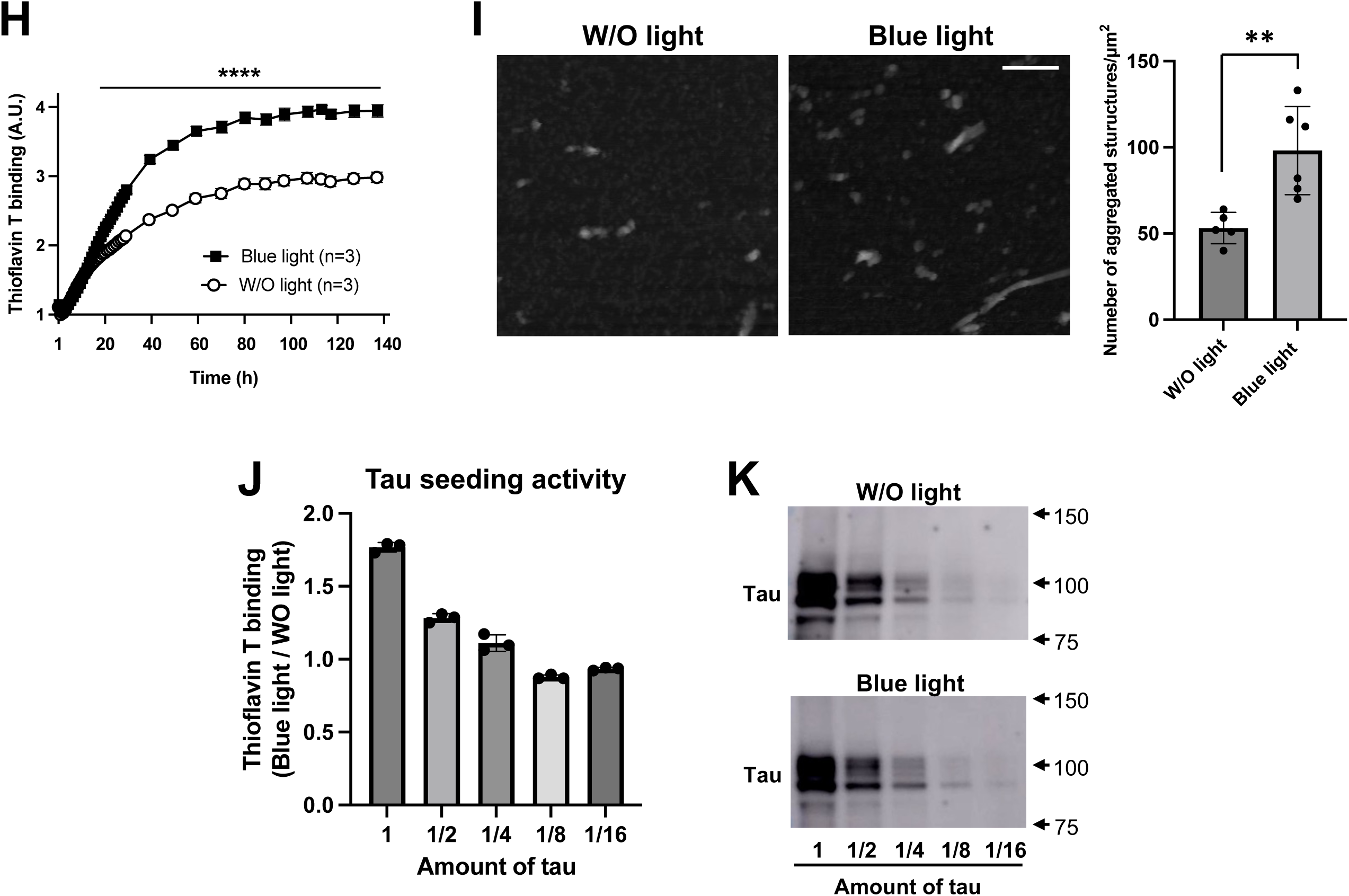
Characterization of solid-like state tau in cells expressing OptoTau-ΔN. (A-E) The cells expressing OptoTau-ΔN were exposed to 10 μM nocodazole and blue light for 22 hrs, and lysed by 1% Triton X-100. After the lysates were separated into 5 fractions by sucrose density gradient centrifugation, tau protein and total proteins in Fractions 1(A), 2(B), 3(C), 4(D) and 5(E) were detected with pan-tau JM antibody (right panel) and ponceau-S staining (middle panel), respectively. Levels of tau were normalized to corresponding ponceau-S staining (lower panel). (F,G) The cells were transfected with OptoTau (F) and OptoTau-ΔN (G). After exposure to blue light with nocodazole for 20 hrs, the cells were stained with SNAP-Cell TMR-Star (magenta), and immunostained with LAMP1 antibody (green). Upon blue light exposure, the major axis of lysosomes, colocalized with or without OptoTau (F) and OptoTau-ΔN (G), was measured using ImageJ. The proportion of larger lysosomes, defined as those with a major axis exceeding 1 μm, was normalized to the total number of lysosomes. (H-K) Tau seeding activity was increased in cell lysates from blue light-exposed OptoTau-ΔN than W/O light. Recombinant tau protein was reacted with cell lysates and then polymerized by heparin. Recombinant tau aggregation was monitored by Thioflavin T binding at indicated times (Thioflavin T binding, F (1, 168) = 5636, P<0.0001; Time, F (41, 168) = 1896, P<0.0001; Interaction, F (41, 168) = 107.9, P<0.0001) (F). AFM detected tau aggregated structure in samples at 140 hrs after polymerization reaction (G, left and middle panels). The aggregated number in the AFM images was counted and quantified (G, right panel). (J) The dose-response of tau seeding activity. The amounts of tau were associated with tau seeding activity. Thioflavin T binding of cell lysates with blue light exposure was divided by that from samples without light exposure. (K) Tau protein in lysates was detected with pan-tau JM antibody. All error bars indicate mean ± SD. ** < 0.01, *** < 0.001 and **** < 0.0001 by Student’s t-test (A-E,G), two-way ANOVA and Šídák’s multiple comparisons test (F) or Fisher’s exact test (G). Fr, Fraction; W/O, without.

Recent studies using phase-separating RNPs, including the use of the CRY2 system, showed that the maturation of condensates into aggregates causes the accumulation of lysosomes ^65,66^. We investigated lysosome accumulation in cells expressing OptoTau (forming droplets) and in cells expressing OptoTau-ΔN (forming insoluble aggregates). Under blue light exposure, OptoTau droplets colocalized with LAMP1, a lysosomal marker, but non-colocalized droplets were also observed (Figs. 5F and S9L). Some of the OptoTau droplets in the LLPS appeared to be migrating to the lysosome. In the cells expressing OptoTau-ΔN, aggregates colocalized with LAMP1 (Figs. 5G and S9M).

Upon blue light exposure, the percentage of lysosomes with the major axis exceeding 1 μm and with OptoTau-colocalized LAMP1 was significantly higher than lysosomes with OptoTau-non-colocalized LAMP1 (Fig. 5G). This enlargement of lysosomal structures was reported under conditions of impaired lysosomal degradation ^65^ ^67^. These results imply that OptoTau-ΔN is successfully delivered to lysosomes, but may induce defects in lysosomal degradation.

It has been reported that phosphorylated high-molecular-weight tau is responsible for tau propagation ^68^, and we observed its increase in the cells co-exposed to blue light and nocodazole (see Fig. 4B and C). To test the promotable effect of the blue light-induced insoluble tau oligomers on tau seeding activity, a seed amplification assay was performed ^69,70^. Recombinant tau aggregation with β-sheet structure was significantly increased by treatment of lysates from blue light exposed cells compared with that obtained from cells not exposed to blue light (Fig. 5H). Atomic force microscopy (AFM) has been used in numerous studies to visualize tau aggregated structures including granular tau oligomers and tau fibrils ^25,26,71,72^. The number of recombinant tau aggregates formed in lysates of blue light exposed cells was significantly higher than that in lysates from cells not exposed to blue light (Fig. 5I). The tau seeding activity was dependent on the amount of tau (Figs. 5J and K). These results indicate that ΔN-P301Ltau-formed oligomers can augment tau seeding activity.

## Discussion

Several cellular models of tau aggregation induced by seeded tau exposure have been developed ^73^ ^74^ ^75^ ^76^ ^77^ ^78^. However, the cellular mechanism and tau aggregation pathway that converts normal tau to seed-active tau are unclear. This study investigated the P301Ltau-mediated aggregation pathway in cultured cells and presents a novel mechanism of aggregation, which includes the formation of intermediate aggregates with seeded activity. First, we showed that OptoTau is clustered in the aggresomes upon exposure to blue light. Second, we observed that OptoTau is scattered in the cytoplasm and forms liquid-phase tau droplets when aggresomes are eliminated using nocodazole. Lastly, we showed that OptoTau-ΔN induces irreversible aggregates including granular tau oligomer in the cytoplasm and confirmed an increase in tau seeding activity. Collectively, our findings support a model whereby P301Ltau clustering is accumulated in aggresomes initially, and disruption of the aggresomal process leads to tau aggregation through LLPS, ultimately forming in tau seed.

### The cellular mechanism of tau droplet formation and its role in neurofibrillary tangles formation

#### Accumulation of tau in aggresomes

Numerous studies using human subjects, animals, and cultured cells have consistently shown the presence of hyperphosphorylated tau in cells during the emergence of tau pathology ^8,79-81^ ^82^. Our results showed that oligomers of OptoTau are in a hyperphosphorylated state, which is consistent with previous reports. Small tau oligomers, such as dimers and trimers, were detected on microtubules ex vivo ^83^. Phosphorylation of tau has been shown to promote tau axonal transport^84,85^. P301Ltau transgenic mice show the ectopic expression of tau by somatodendritic mislocalization ^86^, and an increase in tau phosphorylation at S400 and S422 residues in the C-terminal region ^21^. The results of previous reports and our findings suggest that the oligomer of OptoTau may move retrogradely on microtubules and reach the aggresomes.

The formation of aggresomes as a compensatory change has been considered to occur when the capacity of the proteasome is exceeded by producing aggregation-prone misfolded proteins ^46^. Inhibiting the proteasome drives the formation of tau-positive structures with the morphology of aggresomes ^87,88^. The aggresome marker HDAC6 contributes to the formation of tau aggresomes ^87^. These findings suggest that aggresomes provide an alternative pathway for the degradation and removal of harmful tau aggregates ^89,90 91^. In contrast to previous studies, this study showed that tau oligomers were deposited on the aggresomes in the absence of proteasome inhibitors. This result suggests that the degradation of tau on the aggresomes may occur even if the ubiquitin-proteasome system is not impaired at an early stage. The formation of tau clusters in the tau-CRY2 experiments supports this interpretation. Therefore, our results suggest that the early stages of tau oligomer formation might lead to the degradation of tau-CRY2 through aggresome formation.

#### Transition of liquid phase to solid phase

Our findings show that OptoTau clustered on microtubules and tau phosphorylation increased by blue light exposure. Zhang and colleagues, using a full length wild-tau construct fused to CRY2-mCherry, showed that tau phase separation as a condensate enhanced its binding to microtubules ^92^. The proline-rich domain (a.a.151-245) in tau drives LLPS via the control of its phosphorylation state ^92^, consistent with our findings. Pseudo-phosphorylated tau at S262 (S262E-OptoTau, Fig S6A; S262E-OptoTau-DN, Fig S6B), which strongly reduces the binding of tau to microtubules ^43,93^, was clustered in the cells by blue light exposure. Further, in differentiated Neuro2a cells expressing S262E-OptoTau-DN, granular tau cluster formation (Fig. S6C) and tau phosphorylation (Fig. S6D) were increased by blue light exposure without nocodazole. These results indicate that the clustering and phosphorylation of OptoTau may not be attributed to the artificial effects of nocodazole. Pretangle, the most commonly observed neuronal lesion in AD, is a poorly formed cytoplasmic inclusion, which is positive for phosphorylated tau ^94^. Because the droplets of OptoTau in the cytoplasm are hyperphosphorylated, but not insoluble aggregates, like neurofibrillary tangles (NFTs), the droplets of OptoTau can be considered an intermediate form preceding the formation of tau aggregates. Jiang and colleagues detected the forming tau oligomer by blue light exposure in cultured cells expressing 1N4R-wild type tau with CRY2olig ^95^. Mutant P301Ltau significantly enhances both tau droplet formation^29^ and tau oligomer formation ^96^, compared with wild-type tau in vitro. Recent pathological analysis has shown that the cytoplasmic AT8-positive pretangle before forming NFTs, appears to be in droplet form ^97^. Taking these findings together, in the cytoplasm, microtubule-unbound full length-P301Ltau initially forms droplets through LLPS, leading to the formation of tau oligomers.

Studies have demonstrated that the N-terminal region of tau serves to suppress aggregation ^98^ ^99^. Our findings indicate that clustering of OptoTau-ΔN results in the formation of insoluble granular tau oligomers. It has been established that the microtubule-binding repeat domain of 4R-Tau (K18)^39^ and tau truncated by caspase ^100^ and calpain I ^58^ exhibit a heightened or fast propensity for aggregation in comparison to full length-tau. Ca^2+^ overload and influx by neuronal hyperexcitability may induce tau cleavage ^101^ ^102^. In fact, AMPA and NMDA receptor stimulation increased the local somatodendritic translation and hyperphosphorylation of tau protein ^103^. Additionally, the microtubule-binding repeat (MTBR) undergoes LLPS ^28^. Cryo-EM studies have shown that tau sequences containing amino acid residues 297-391 of tau (referred to as dGAE) are the common core structure of fibrils in tauopathies ^11-16^. Moreover, the dGAE sequence was found to be capable of inducing the formation of AD-like fibrillar core structures in vitro ^104^. RNA bound the C-terminal regions of tau at R406 and H407, enhance the integrity of aggregated tau ^105^. Therefore, our results show that ΔN-P301Ltau forms insoluble aggregates that are consistent with findings of previous studies.

A portion of the OptoTau droplets formed by LLPS was observed migrating to the lysosome. Upon exposure to blue light, OptoTau was colocalized with p62, a ubiquitin receptor that facilitates the clearance of ubiquitin-tagged protein aggregates through autophagy ^106,107^. Misfolded oligomers are transported to the aggresomes and removed from active sites ^91^. Given our findings and previous reports, it is likely that oligomers derived from OptoTau droplets are transferred to lysosomes for degradation. The fragmentation of OptoTau, OptoTau-ΔN, was delivered to lysosomes and associated with an increase in larger lysosomes, suggesting that its insoluble aggregates may impair the autophagy process. Tau accumulation has been shown to induce autophagy deficits ^108^. In an AD model mouse, the size of autolysosomes was increased ^109^. In a human study, Nixon and colleagues first reported direct evidence linking autophagy to AD ^110^. Further studies indicate that attenuation of the autophagy pathway is a risk factor for age-related neurodegenerative disease, including AD ^111^ ^112^ ^113^. Therefore, the deposition of OptoTau-ΔN might attenuate the endo-lysosomal pathway contributing to AD.

Tau seeds play an important role in tau propagation ^68,114-117^. We found that the tau oligomers observed in OptoTau-ΔN-expressing cells have tau seeding activity. Considering this, in combination with the fact that thioflavin T is responsible for β-sheet rich fibrils structure ^25^, N-terminal cleaved tau is suggested to be the tau seed to form fibrils. Prior to the formation of seeded tau, P301Ltau oligomers accumulated along microtubules and in the aggresomes. Pathological analysis of human brains aged 1 to 100 years shows that the earliest change is AT8-positive pretangle in proximal axons of the locus coeruleus, which precedes AD tangles ^118^. The events observed in our cells mimic such human tau pathology. Although the question of the nature of seed tau needs careful study, dextran tau forms granular tau oligomer but not long fibrils (Fig. S7), and the dextran-formed tau aggregates enhance the propagation of endogenous tau in C57BL/6J mice ^119^. Aggregates of dGAE increase endogenous tau phosphorylation ^120^, and phosphorylated high-molecular-weight tau derived from AD brain shows tau seeding activity ^68^. These results suggest that the granular tau oligomers generated in this study have seed activity.

### Novelty and usefulness of the light-induced model of tau aggregation using CRY2olig

Previous studies have shown that elucidating the initiation of the granular tau aggregation process is essential to better understand tauopathies ^25,26,121^. In this study, we found that OptoTau-ΔN accumulates in the cytoplasm and forms granular tau oligomers. While previous research has shown that recombinant full length-P301Ltau can form granular tau oligomers in vitro^96^, our study is the first to produce granular tau oligomers in cells by optogenetic manipulation. In addition, for the first time, we demonstrate the intracellular formation of granular tau oligomers with seeding activity. These results imply that P301Ltau is converted from a liquid phase with soluble properties to a solid phase with insoluble properties in cells acquiring seed activity. In the liquid phase, tau local concentration is induced by aggresome disruption and LLPS, which precedes aggregation. Thus, this study presents a novel mechanism for the aggregation pathway leading to the formation of intermediate aggregates with seed activity. These intracellular events may be therapeutic targets in tau-related diseases to inhibit the formation of granular tau oligomers prior to forming NFTs. The circularity of tau condensates decreased in cells expressing OptoTau-ΔN and exposed to blue light, but this did not lead to tau fibril formation. We speculate that the lack of fibril formation is due to the low amount of granular tau oligomers in the cells. Because granular tau oligomers self-associate to form fibrils ^25^, tau fibrils may be formed when the amounts of granular tau oligomers reach the threshold point.

### Limitations of the study and future directions

This study, using CRY2olig fusion tau protein, cannot exclude the possibility of effects of CRY2olig or blue light exposure on other cellular processes. We observed the aggregation of P301Ltau in Neuro2a cells, an immortalized cell line. However, the formation of tau aggregation may differ between Neuro2a cells, other cell lines, and cells in living organisms. Further analysis using primary neurons or transgenic mice expressing tau with CRY2olig may provide a more precise understanding of the biological mechanism of tau aggregation. Although tau pathology is observed in both neurons and glial cells, the aggregation of tau in glial cells has not been explored in this study. This CRY2olig system might be useful in investigating the mechanism of tau aggregation in glial cells, where the contribution of microtubule-organizing centers in the formation of aggresomes is well-established ^46^. Granular tau oligomers have been reported to be associated with cell death ^21,122^, but this study did not address cell death, as the immortalized Neuro2a cell line has an intrinsic proliferative capacity that may mask tau-induced cell death. The precise mechanism by which granular tau oligomers form in the cytoplasm remains unclear. Investigating the mechanism of tau aggregation in cells could lead to the identification of new targets for tau-related drug discovery.

## Conclusion

In conclusion, our study proposes a new model of cellular tau aggregation in that P301Ltau fused with CRY2olig forms soluble tau oligomers in the cytoplasm by LLPS. Subsequently, N-terminal cleaved soluble oligomer forms insoluble granular tau oligomers, which have seeding activity. Further investigation into the mechanisms of this model may provide insights for the development of novel therapies for tauopathies.

## Acknowledgments

This work was supported by a Grant-in-Aid for Scientific Research on Innovative Areas ‘Singularity Biology (No. 8007)’ (18H05414 and 18H05410) of MEXT to H.B. and H.Y., respectively; JSPS KAKENHI Grant Number 20K06896 and 23K05993 to Y.S., 23K18116 to H.B., and 22K15399, 22H05574 and 24H01327 to I.S.; Takeda Science Foundation to Y.S.; Abe Yoshishige Foundation to Y.S.; and AMED under grant number JP21wm0425016 to A.T. Part of the English language refinement in this paper was supported by ChatGPT. We greatly appreciate the support of the NAI, Inc. (Yokohama, JAPAN) for the English language review.

## Author contributions

Y.S. conceived and designed the study and drafted this article. H.Y. constructed cDNA. R.K. critically supported drafting the article. I.S. immunostained mitochondria and lysosomes. Y.S, H.B., and A.T. critically discussed the design of the experiments and the article.

## Declaration of interests

The authors declare no competing interests.

## STAR METHODS

### KEY RESOURCES TABLE

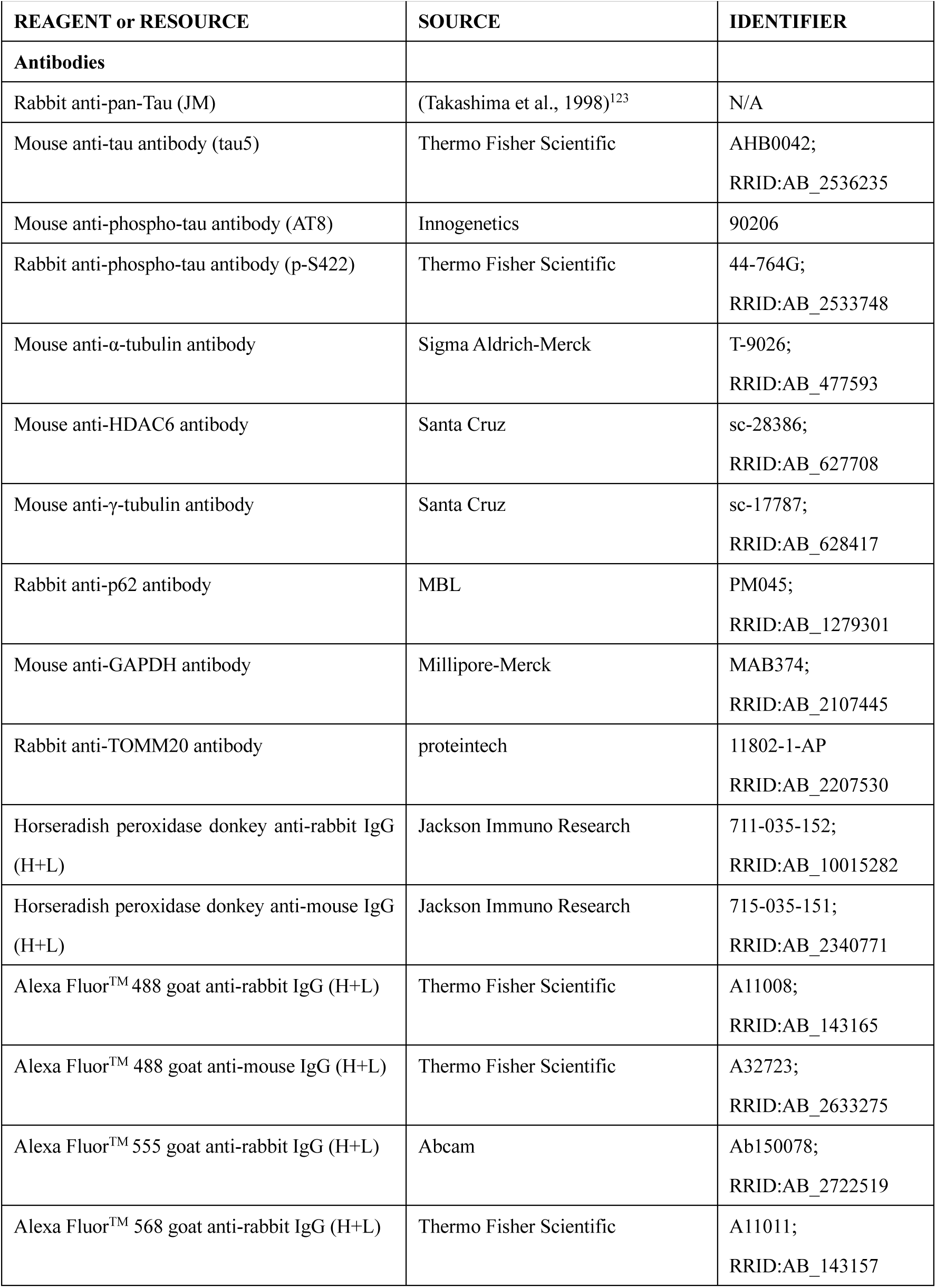
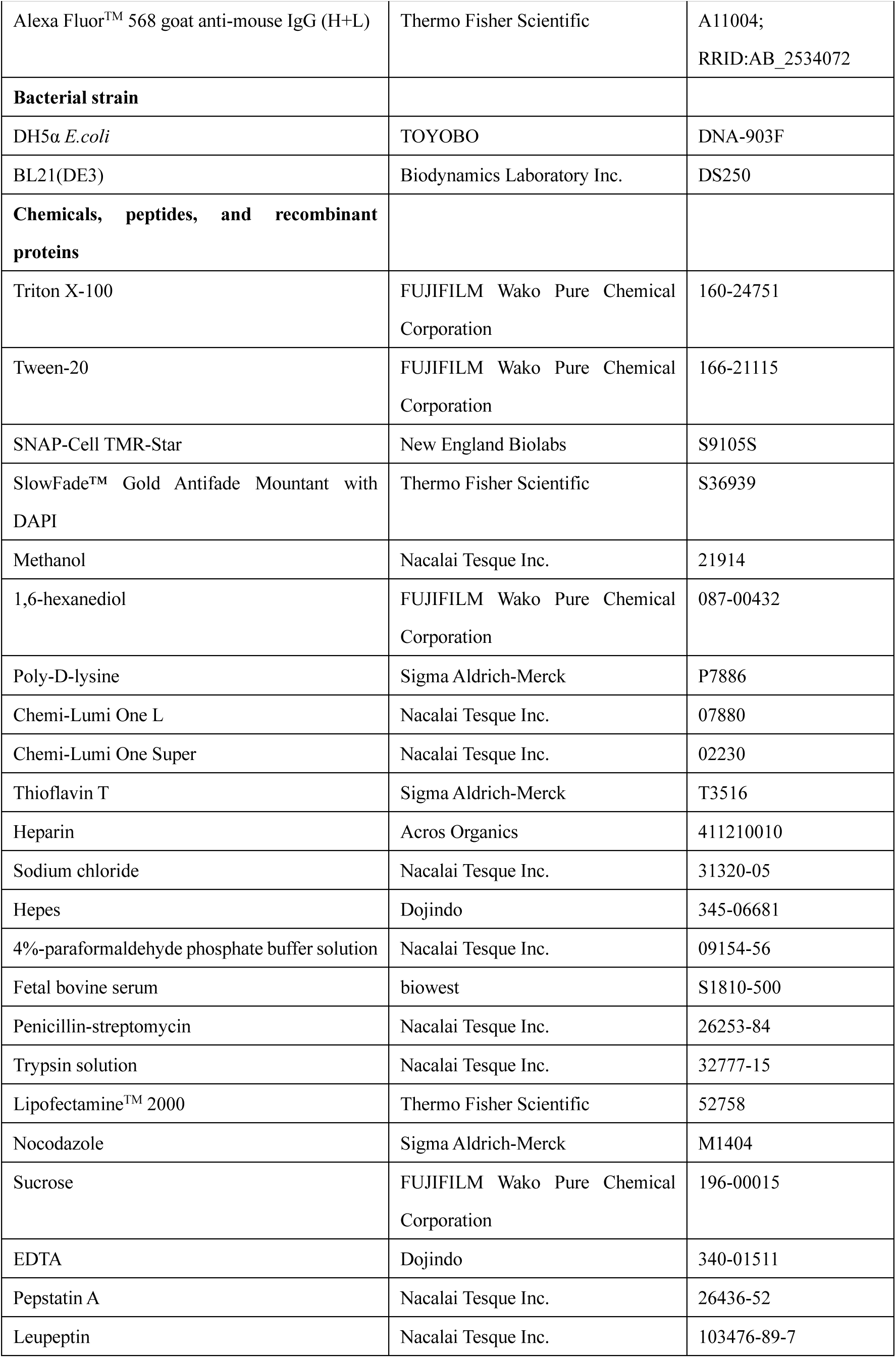
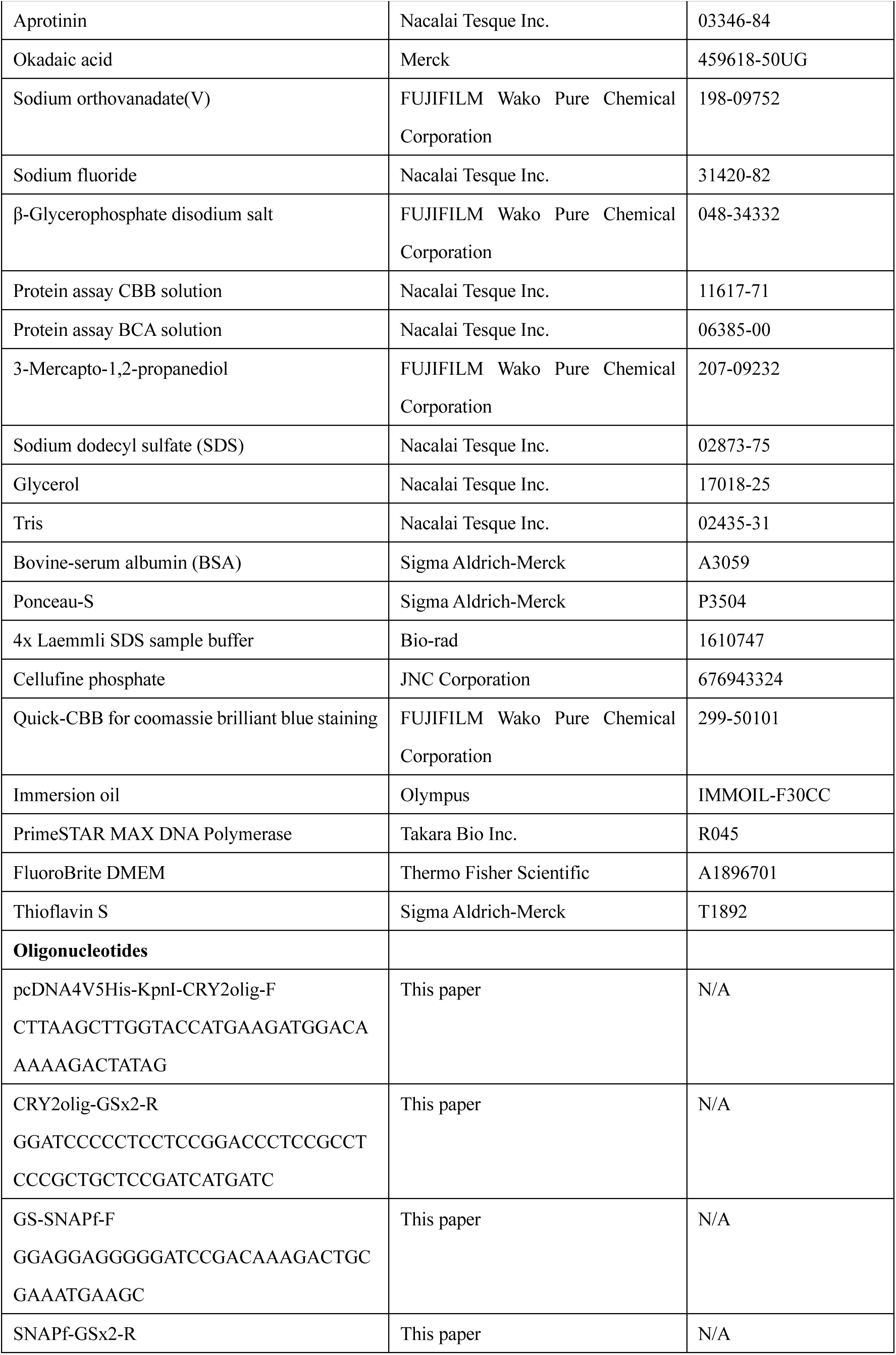
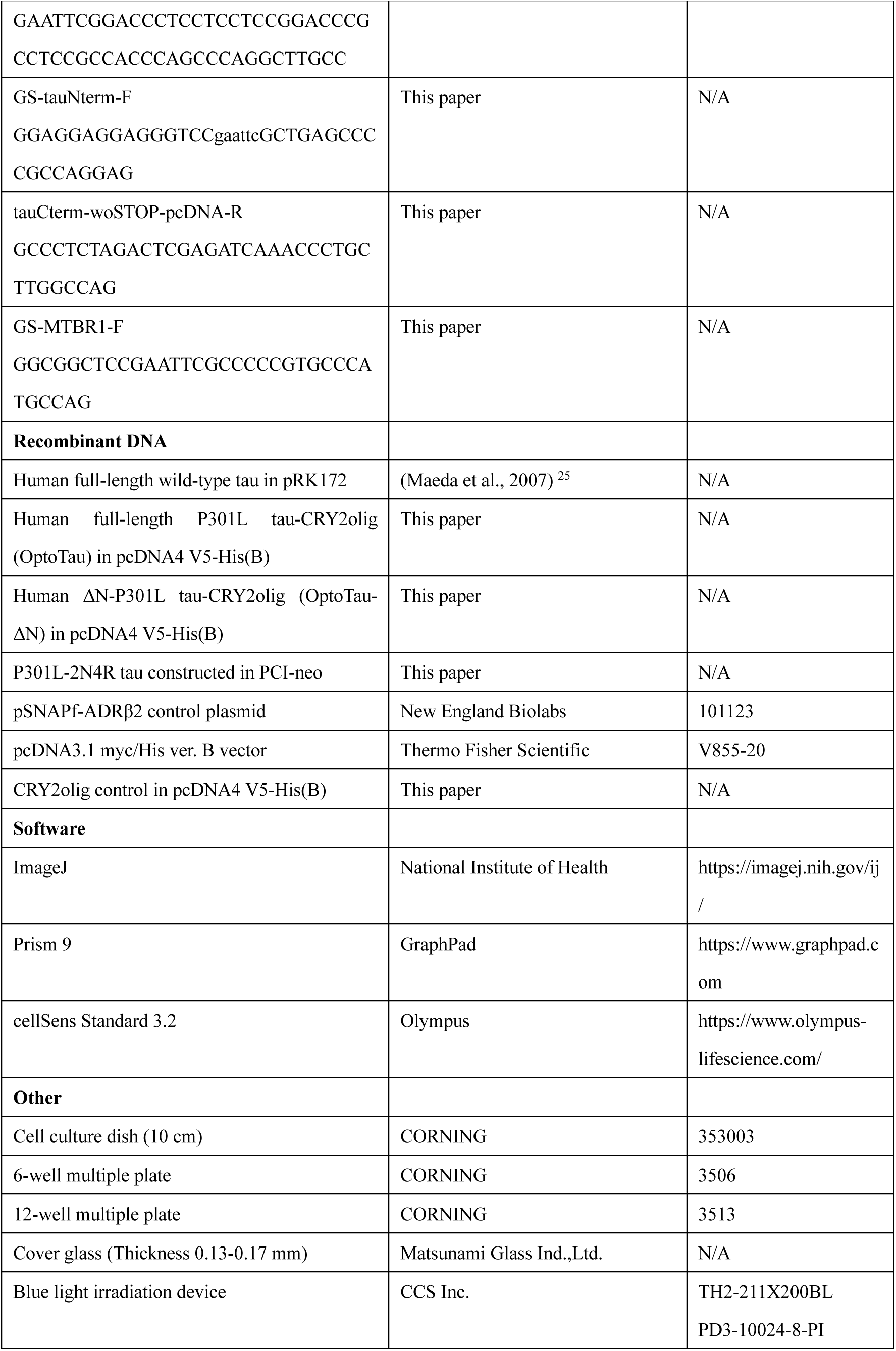
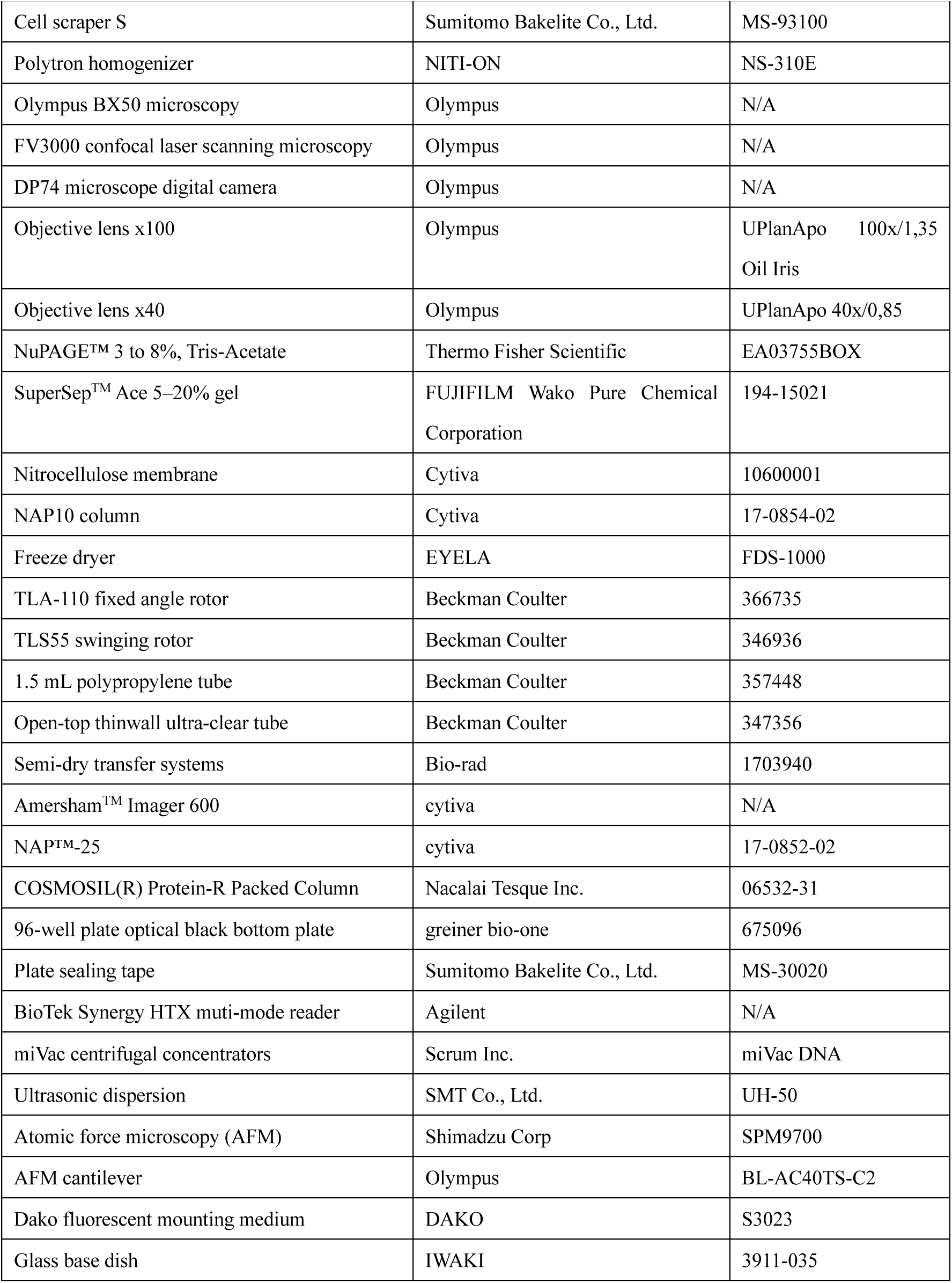

### RESOURCE AVAILABILITY

#### Lead contact

Further information and requests for resources and reagents should be directed to and will be fulfilled by the lead contact, Yoshiyuki Soeda.

#### Materials availability

All unique/stable reagents generated in this study are available from the lead contact with a completed materials transfer agreement.

#### Data and code availability

- All data reported in this paper will be shared by the lead contact upon request.
- This paper reports no original code.
- Any additional information required to reanalyze the data reported in this paper is available from the lead contact upon request.

## EXPERIMENTAL MODEL AND SUBJECT DETAILS

### Cell line

The mouse neuroblastoma cell line, Neuro2a, was cultured in Dulbecco’s Modified Eagle’s medium supplemented with 10% fetal bovine serum (FBS) and 1x penicillin-streptomycin. The culture was maintained in a 10 cm dish at 37 °C under 5% CO2.

## METHOD DETAILS

### cDNA construction

The cDNAs of tau and SNAPf were amplified using PrimeSTAR MAX DNA Polymerase from P301L-2N4R tau constructed in PCI-neo and pSNAPf-ADRβ2 control plasmid, respectively. The CRY2olig cDNA was prepared through site-directed mutagenesis (E409G) of CRY2 gene that was obtained from Addgene. The PCR products were subcloned into pcDNA3.1 myc/His ver. B vector. All primers used for this study are listed in the KEY RESOURCES TABLE. The cDNAs were transformed into DH5α, and their glycerol stocks were stored at −80 °C.

### Transfection

Neuro2a cells at about 90% confluency were detached using 0.25 g/L trypsin and counted.

For biochemical analysis, cells at 2.0 x 10^5^ cells /ml were plated in a 6-well plate (2 ml) or 12-well plate (1 ml). For immunofluorescence analysis, cells at 1.0 x 10^5^ cells /ml were plated onto cover glass coated with poly-D-lysine. cDNAs (1 μg) were transfected by lipofectamine 2000 according to the instruction manual 1 day after culturing. The transfection reagents were removed by a medium change 6-8 hrs after transfection.

### Irradiation of blue light

The transfected cells were exposed without or with 10 μM nocodazole in serum-free DMEM 1 day after transfection. The cells were placed onto a transparent plastic case with a height of 3 cm ^124^. After setting the scale of the blue light irradiation device to 029, the cells were irradiated with blue light at 0.87q0.16 mW/cm^2^ for 1-24 hrs. To serve as a control, aluminum foil was utilized to enclose the cell plates and prevent exposure to ambient light.

### Neuro2a cell differentiation and blue light irradiation

After removal of the transfection reagent, the cells were cultured in DMEM containing 1% FBS and 20 μM retinoic acid for 2 days. Subsequently, the cells were exposed to blue light with nocodazole in DMEM containing 1% FBS and 20 μM retinoic acid for 1 hr.

### Preparation of cell homogenate

The cells in the medium were scraped, and then cell samples including the medium were centrifuged (15000 rpm, 2 minutes, 4r). After removing the supernatants, the pellets were homogenized in a buffer (250 mM sucrose, 1 mM EDTA, 10 mM Hepes [pH=7.4]) containing protease inhibitors (5 μg/ml pepstatin A, 5 μg/ml, leupeptin, 2 μg/ml, aprotinin), and phosphatase inhibitors (1 μM okadaic acid, 1 mM sodium orthovanadate(V), 1 mM sodium fluoride, and 1 mM β-glycerophosphate disodium salt) using a polytron homogenizer for 30 seconds at a setting to 10. After the homogenized solutions were centrifuged (3000xg, 10 minutes, 4r), supernatants were recovered^125^. The protein concentration of the samples was measured with the Bradford assay. To analyze under reducing condition, samples were suspended in 2.5% 3-mercapto-1,2-propanediol-containing 1x Laemmli SDS sample buffer followed by boiling for 10 minutes. Under non-reducing condition, the samples were suspended in sample buffer containing 0.5% SDS, 10% glycerol, 0.002% bromophenol blue and 62.5 mM Tris-HCl (pH=6.8).

### Staining of tau with CRY2olig

The tau cDNAs were fused to CRY2olig with a SNAPf tag (figure S1), which can covalently bind to chemical fluorescent dyes. SNAP-Cell TMR-Star (excitation wavelength, 554 nm; fluorescence wavelength, 580 nm) was used to label the tau protein with CRY2olig in cells. The cells on cover glass coated with poly-D-lysine were fixed with prewarmed 4% paraformaldehyde (PFA) for 15 minutes at room temperature. After washing with phosphate-buffered saline (PBS), the cells were treated with 0.1 μM SNAP-Cell TMR-Star and 1% bovine serum albumin (BSA) in PBS, and incubated at 4r in a moist and dark box over-night.

### Thioflavin S staining

The cells were transfected with cDNA of OptoTau-ΔN and the CRY2oligo vector control, and subsequently exposed to blue light with 10 μM nocodazole for 18 hrs. After exposure, the cells were fixed with prewarmed 4% paraformaldehyde (PFA) for 15 minutes and permeabilized with 0.1% Triton X-100 for 3 minutes at room temperature. The cells were then stained with 1.57 μM thioflavin S in PBS buffer for 40 minutes at room temperature.

### Immunofluorescence

The cells were fixed with prewarmed 4% PFA for 15 minutes at room temperature to detect HDAC6, p62, and TOMM20. To detect α-tubulin and γ-tubulin, the cells were fixed by methanol at −30r for 15 minutes followed by 4% PFA at room temperature for 5 minutes. After permeabilization with 0.1% TritonX-100, the cells were blocked by 1% BSA containing 0.1% Tween 20 in PBS buffer. The cells were treated with primary antibodies (anti-α-tubulin antibody [1:500], anti-HDAC6 antibody [1:100], anti-γ-tubulin antibody [1:100], anti-p62 antibody [1:100], and TOMM20 [1:100]) and SNAP-Cell TMR-Star (0.1 μM) in 1% BSA containing 0.1% Tween 20 in PBS buffer and incubated at 4r over-night. After washing with PBS, incubation at room temperature for 0.5 ∼ 1 hr was conducted in 2^nd^ antibodies with Alexa fluor (1:1000) in 1% BSA containing 0.1% Tween 20 in PBS buffer.

### Microscopy observation and image analysis

After staining, the cells were mounted using SlowFade^TM^ gold with DAPI or, Dako fluorescent mounting medium and observed under reflected light microscopy (figures, 3A, 3B, 4G, 4H, 4I, 4J and 4K) and confocal microscopy (figures 1D, 1F, 2E, 2F, 4D, 4F, 4L, 5F and 5G) with oil immersion objective lenses x100 or x40. Images were obtained using a digital camera DP74 and image software, cellSens Standard 3.2. Tau clusters stained by SNAP-Cell TMR-Star were counted. To quantify tau intensity (Fig. 1D-d, 2E-d and 4D-d), during blue light exposure, ROIs were delineated within tau clusters using a freehand tool. In the absence of light irradiation, ROIs were manually defined to correspond to the area of the perinuclear region that is exposed to blue light (Fig. S11). Levels of tau clustering were normalized to corresponding total tau, represented by the whole-cell magenta fluorescence intensity. To quantify the thioflavin S signal under blue light exposure (Fig. 4F), images of SNAP-Cell TMR-Star-positive magenta fluorescence and thioflavin S-positive green fluorescence were merged. The number of clusters bearing merged signals was normalized to the corresponding clusters bearing magenta signals. To measure the circularity of tau clusters in cells (Fig. 4I), analyze particles in ImageJ was performed after images were adjusted for brightness/contrast and binarized. Circularity was calculated using the formula 4π multiplied by the area divided by the square of the perimeter. A circularity value of 1.0 signifies a perfect circle, while values nearing 0.0 indicate a progressively elongated shape ^126^. To quantify the size of lysosomes under blue light exposure, LAMP1-positive green fluorescence was adjusted by threshold, binarized and counted. The major axis of each lysosome was measured using the analyze particles in ImageJ. Lysosomes were categorized based on whether they colocalized with tau or not, and each category was analyzed separately. The ratio of larger lysosomes with the major axis exceeding 1 μm to all lysosomes was calculated.

### Experiments of light off and 1,6-hexanediol exposure

After cells expressing OptoTau and OptoTau-ΔN were irradiated with blue light for 24 hrs, the cells were cultured in DMEM containing 10 μM nocodazole without FBS. For the turning light off experiment, the culture was performed for 18 hrs without blue light irradiation. The cultures were carried out under aluminum foil to prevent the influence of ambient light. For the 1,6-hexanediol exposure experiment, cells were cultured in DMEM with 4% 1,6-hexanediol for 10-60 minutes at 37 °C in a CO_2_ incubator. Tau proteins in these cells were stained with SNAP-Cell TMR-Star.

### Preparation of sarkosyl-insoluble fraction

Cells in three wells of the 6-well plate were collected 24 hrs after blue light irradiation. A negative control is cells expressing an empty vector. To prepare the positive control, recombinant tau (10 μM) was polymerized with heparin for 7 days, and diluted to 1 μM. Cells and positive control were homogenized in 300 μl of the homogenization buffer (see Preparation of cell homogenate), and centrifuged at 1000 x g for 10 minutes to remove nuclear. One hundred eighty μl of the supernatant (total fraction) was transferred into 1.5-ml tube, and mixed 20 μl of 10% TritonX-100 (final concentration, 1%). After rotating for 20 minutes at 4 °C to solubilize other organelles, the supernatants were centrifuged (65,000 rpm, 1 h, 4 °C) in a TLA110 swinging rotor. After removing the supernatant, 200 μl of 1% sarkosyl was added to the pellets. The pellets were sonicated for 1 second at a setting to 1. This operation was repeated until the pellet was invisible to the human eye. After rotating for 1 hr at 37 °C, centrifugation (65,000 rpm, 1 hr, 4 °C) was conducted. Pellets were washed with milliQ water to remove trace soluble tau, and solubilized in 70% formic acid. After the formic acid in the samples was removed by evaporating, pellets were suspended in 30 μl of 1x Laemmli SDS sample buffer containing 2.5% 3-mercapto-1,2-propanediol and boiled. If the color of the sample was yellow, it was restored to blue by adding a small amount of 1 M Tris (pH=7.4).

### Sucrose density gradient centrifugation

Sucrose density gradient centrifugation was performed as previously described ^26^ with minor modifications. Cell homogenates from OptoTau-ΔN-expressing cells were solubilized with 1% TritonX-100 at 4 °C for 20 minutes to solubilize cell organelles ^127^. High-molecular-weight tau formed by irradiation of blue light was not solubilized by 1% Triton X-100 (figure S8). Sucrose step gradients (each containing 450 μl of 10, 20, 30, 40 and 50% sucrose in buffer including 10 mM hepes and 1 mM EDTA) were prepared in an open-top thinwall ultra-clear tube. The samples (450 μl) were layered on top of the sucrose step gradients, and centrifuged (50,000 rpm, 2 hrs, 4 °C) in a TLS55 swinging rotor and separated into 5 fractions. The recovered fractions were suspended in the non-reducing sample buffer and applied to SDS-PAGE western blotting.

### SDS-PAGE western blotting

The samples from the preparation of cell homogenate and the sucrose density gradient centrifugation were separated by NuPAGE™ 3 to 8% tris-acetate gel or SuperSep^TM^ Ace 5–20% gel. Proteins in the gels were transferred onto nitrocellulose membranes using semi-dry transfer systems. The membrane was blocked with 3% milk in PBS-T for 1 h at room temperature. They were probed with antibodies including JM (1:1000-1:2000), AT8 (1:750), p-S422 (1:1000) and GAPDH (1:5000) antibodies in 3% BSA for overnight at 4 °C. After washing the membrane with PBS-T, blots were incubated with horseradish peroxidase-linked secondary antibodies (1:5000) in PBS-T. The membranes were washed and examined by enhanced chemiluminescence detection (Chemi-Lumi One L or Chemi-Lumi One Super) on an Amersham^TM^ Imager 600. To quantify band intensity, ROIs in the image were defined by a rectangle, and the gray mean value was measured using ImageJ. Levels of phospho-tau were normalized to corresponding total tau detected by the JM antibody.

### Preparation of recombinant tau protein

Human wild-type tau (2N4R) cDNA was cloned into a pRK172 vector^25^. Recombinant tau was expressed in *E. coli* BL21 (DE3) and purified by the modified method previously reported ^25^. After sonication and boiling of the *E. coli*, recombinant tau protein in the heat-stable fraction was purified by ion-exchange chromatography (Cellufine Phosphate), ammonium sulfate fractionation, gel filtration chromatography (NAP10 column) and reverse phase-HPLC (COSMOSIL Protein-R Waters). After freeze-drying, recombinant tau proteins were dissolved in milliQ water and stored at −80 °C as a stock solution. The concentration of tau protein was determined relative to BSA as the standard by coomassie brilliant blue staining (Quick-CBB) after the separation of tau protein by SDS-PAGE.

### Tau seeding activity

Cell homogenates were obtained from cells expressing OptoTau-ΔN with or without irradiation of blue light by the aforementioned preparation of cell homogenate. They were incubated in 1% TritonX-100 at 4 °C for 20 minutes to prepare cell lysates. The concentration of total proteins in the cell lysates was adjusted to 0.8 mg/ml. After mixing recombinant wild type-tau protein (10 μM) and Thioflavin T (10 μM) in Hepes buffer (10 mM Hepes, pH=7.4; 100 mM NaCl), 4 μl of the cell lysates were added. The samples were incubated at room temperature for 15 minutes, and then heparin (0.06 mg/ml) was added. Three reactions were prepared in a 96-well plate optical black bottom plate to a final total volume of 25 μl. The plate was covered with sealing tape and incubated at 37 °C in a plate reader. The kinetics of tau polymerization were monitored in real-time by measuring the fluorescence intensity every hour at 420/27 nm excitation and 500/27 nm emission from the bottom of the plate. Shaking was performed for 10 seconds of linear shaking at 550 rpm before the reading of fluorescence intensity. For the dose-response analysis of tau seeding activity, the thioflavin T binding of cell lysates exposed to blue light was normalized by dividing it by the average of samples exposed without light.

### Atomic force microscopy (AFM)

The AFM equipped with a dynamic mode was employed for the observations of recombinant tau aggregated structure ^25,128^. After tau seeding activity, 2 μl of tau aggregation mixtures was diluted by 18 μl of milliQ-water and was deposited onto freshly cleaved mica. The mica was incubated for 10 minutes at room temperature in a moist box, and washed by 100 or more μl of milliQ-water. An AFM microcantilever scanned the mica at 0.3 Hz. The resolution of images is 512*512 pixels. After the Images were adjusted by threshold and binarized, the number of aggregated structures was counted by the analyze particles in ImageJ. The threshold for the analysis was set at 2.257 nm on the major axis, which is difficult to distinguish from noise.

### Statistical analysis

All statistical tests were conducted using GraphPad Prism 9. The significance of differences between two groups was assessed by Student’s or Welch’s t-tests following an F-test. Differences between multiple groups were assessed by one-way analysis of variance and Dunnett’s multiple comparisons test or Tukey’s multiple comparisons test. Data from tau seeding activity were analyzed using a two-way ANOVA. The ratio of larger lysosomes was analyzed using Fisher’s exact test. A p-value less than 0.05 was considered statistically significant.

## Notes

### Competing Interest Statement

The authors have declared no competing interest.

### Summary of Updates

Figures 1D, 1E, 1F, 2E, 2F, 4D, 4F, 4L, 4M, 5F, 5G, 5J and 5K are revised. Author list is updated. Page 10 in discussion section is revised. We adopted a color-blind friendly style, substituting red with magenta in all figures. Supplemental files are updated.

