## Supplemental Information for "Intracellular Tau Fragment Droplets Serve as Seeds for Tau Fibrils"

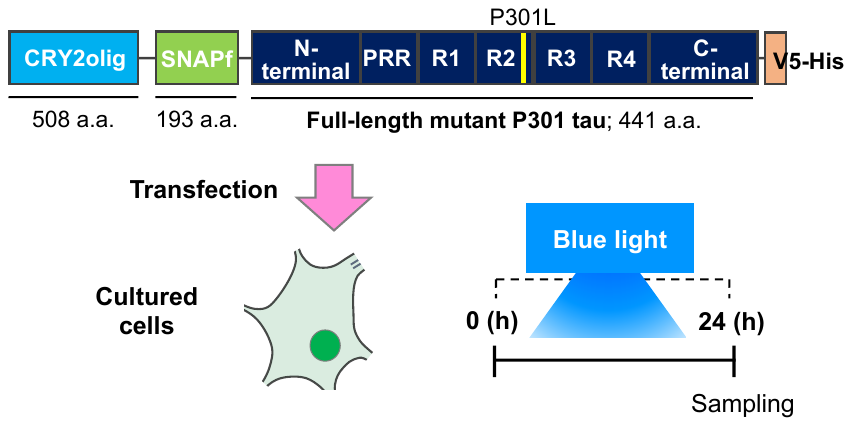

**Figure S1**

**Figure S1. Sequence of OptoTau and experimental designs.**

The human full-length (2N4R isoform) mutant P301Ltau and mutant E490G CRY2olig were fused through a linker, SNAP-tag, which forms a covalent bond with benzylguanine derivatives. The cDNA plasmids were transfected into cultured Neuro2a cells, and irradiated with blue light for 24 hrs.

**Figure S2**

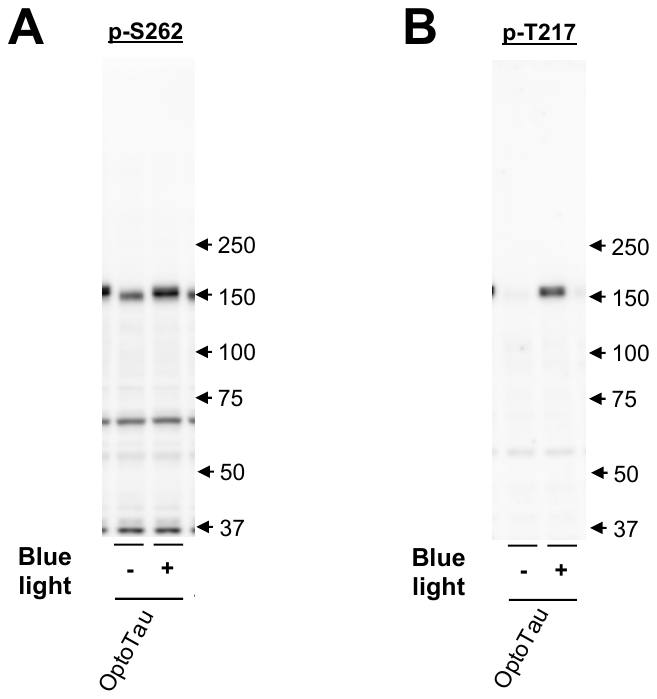

**Figure S2. Induction of tau phosphorylation by blue light in Neuro2a cells expressing OptoTau.**

Tau phosphorylation was detected by p-S262-tau antibody (A) (Thermo Fisher Scientific, 44-750G) and p-T217-tau antibody (B) (Thermo Fisher Scientific, 44-744) in Neuro2a cells expressing OptoTau with or without irradiation of blue light for 24 hrs. p-, phosphorylated.

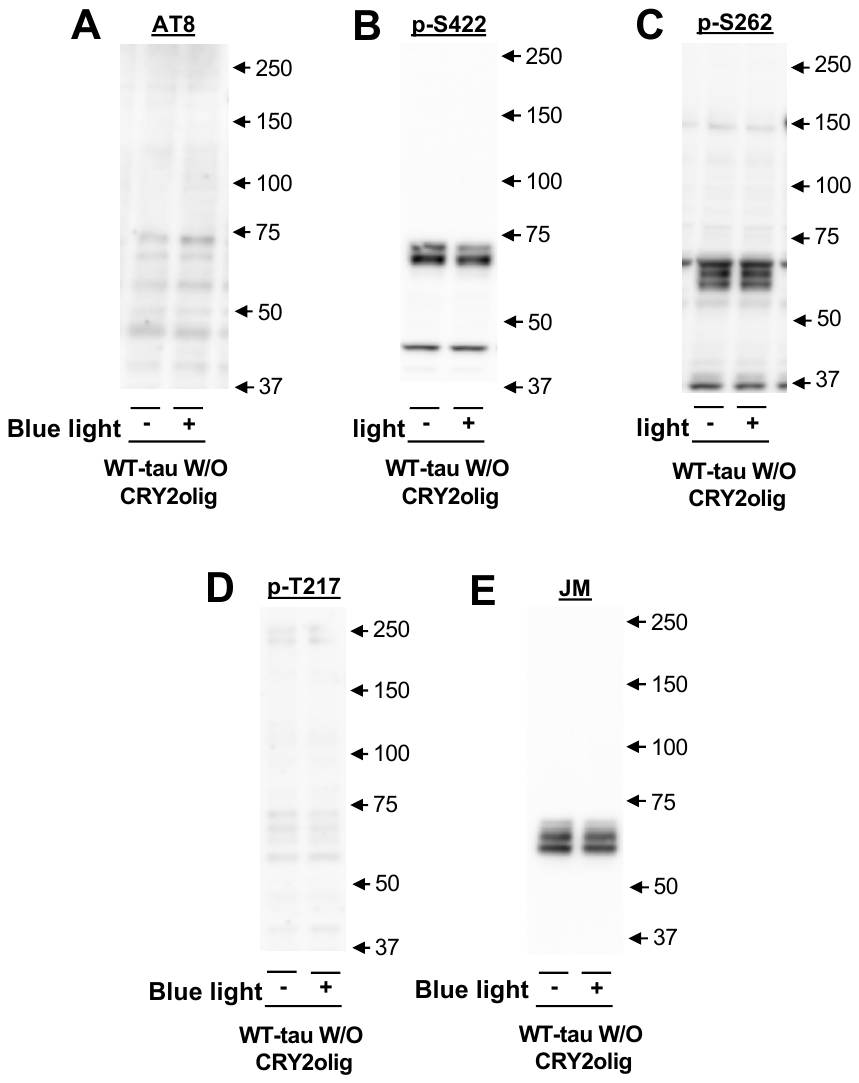

**Figure S3**

**Figure S3. Irradiation of blue light does not induce tau phosphorylation in cells expressing full-length tau without CRY2olig.**

(A-D) Tau phosphorylation was detected by AT8 antibody which recognizes tau phosphorylation at S202 and T205 (A), p-S422-tau antibody (B), p-S262-tau antibody (C) and p-T217-tau antibody (D) in Neuro2a cells expressing wild-type (WT) full-length tau without CRY2olig after irradiation of blue light for 24 hrs.

(E) Tau protein was detected by pan-tau JM antibody in the cells.

p-, phosphorylated.

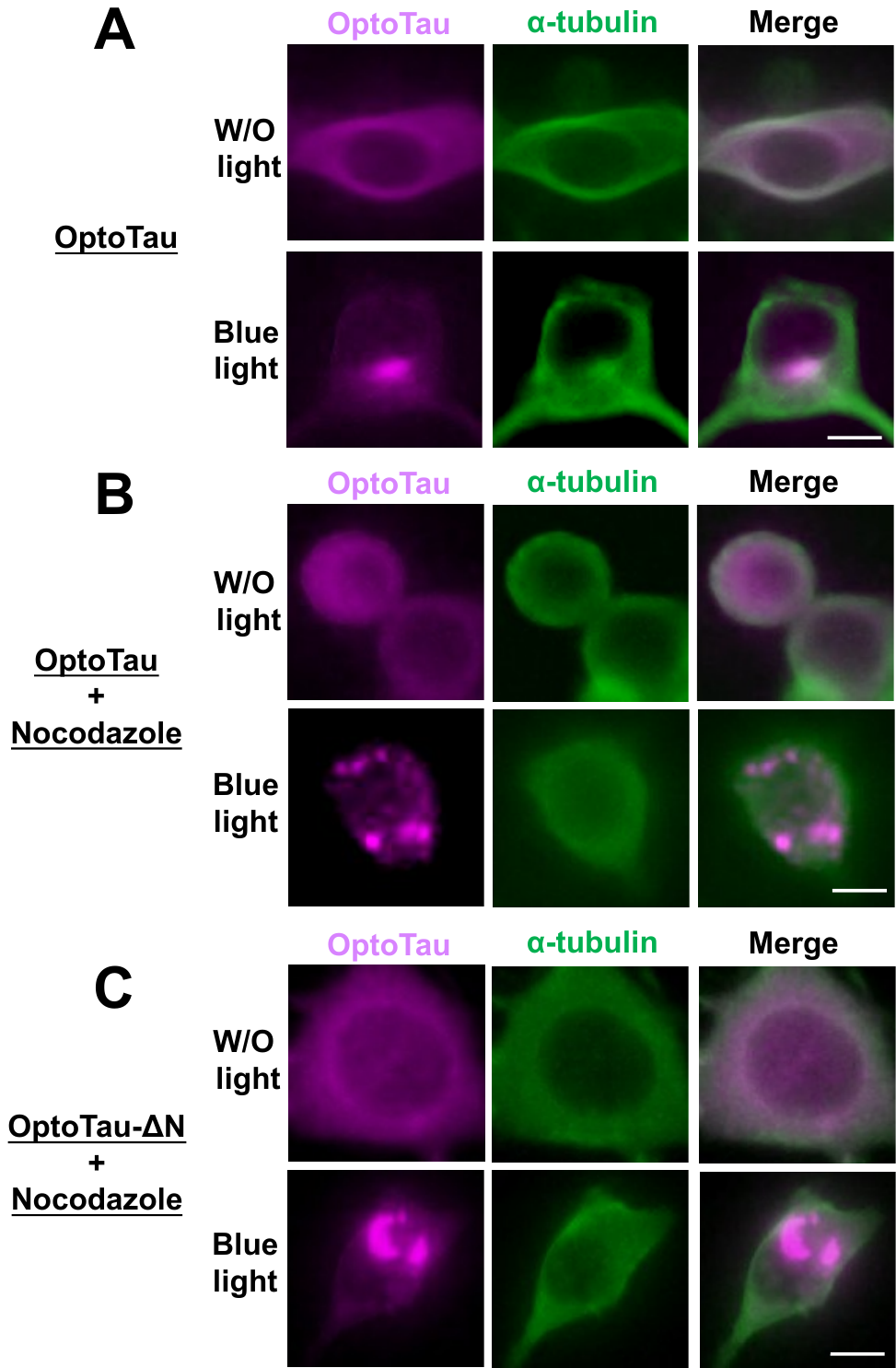

**Figure S4**

**Figure S4. Images of tau clustering were detected with reflected light microscopy.**

Neuro2a cells were transfected with OptoTau (A,B) and OptoTau-ΔN (C), and were exposed to blue light with (B,C) or without (A) 10 μM nocodazole for 20 hrs. The cells were fixed with methanol (15 minutes, -20 ℃) and 4% paraformaldehyde (5 minutes, room temperature). To analyze the colocalization of tau in microtubules by reflected light microscopy (EVIDENT, BX50), the cells were stained with SNAP-Cell TMR-Star to detect tau (magenta) (left panel), and immuno-stained with α-tubulin antibody (green) (middle panel). Images of tau and α-tubulin were merged (right panel). W/O, without. The white bar is 10 μm.

**Figure S5**

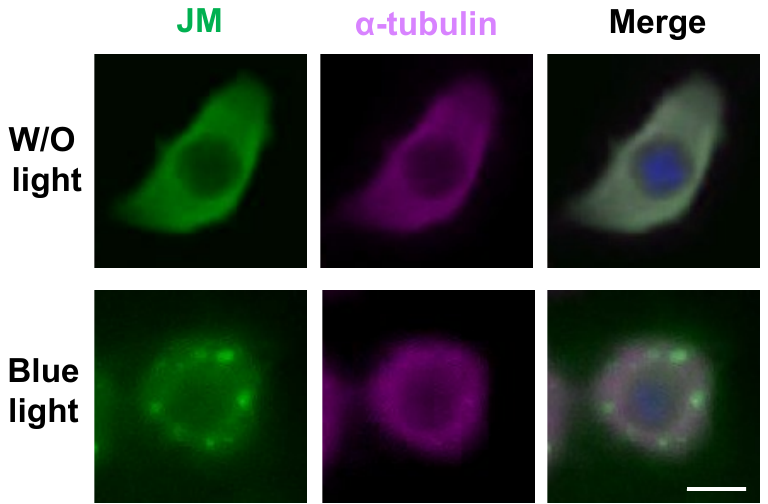

**Figure S5. Tau antibody also detects cytosolic clustering OptoTau under inhibition of microtubule assembly.**

The cells were co-exposed to nocodazole and blue light for 24 hrs, and were immuno-stained with pan-tau antibody (JM) to detect OptoTau (green) (left panel), and with α-tubulin antibody to detect microtubule (magenta) (middle panel). The cells were observed under reflected light microscopy (EVIDENT, BX50). Images were merged (F, right panel). W/O, without. The white bar is 10 μm.

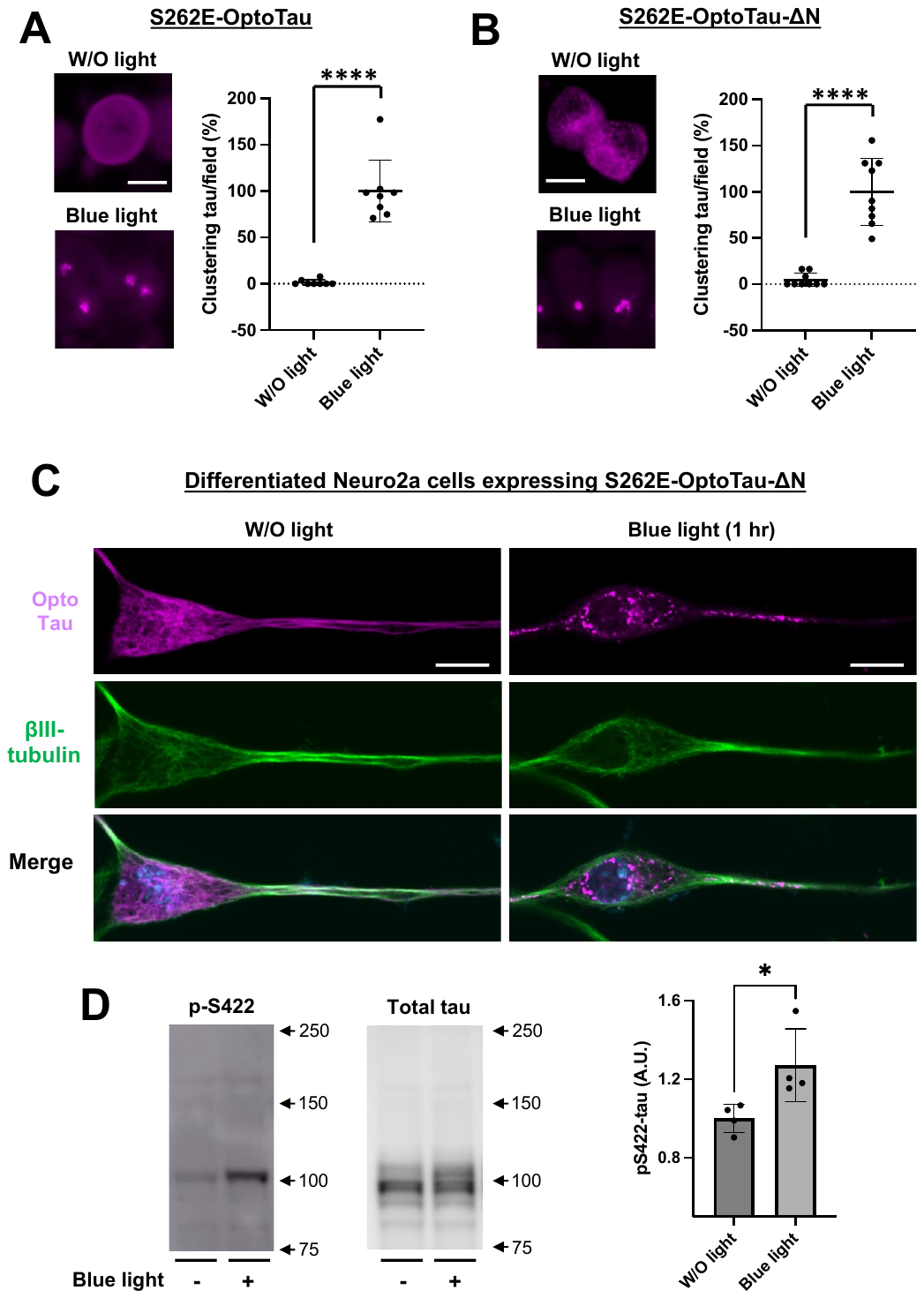

**Figure S6**

**Figure S6. Pseudophosphorylated tau at S262 with CRY2olig is clustered by blue light without nocodazole treatment.**

(A,B) Pseudophosphorylated S262E with full-length (FL)-P301Ltau (S262E-OptoTau; A) and ΔN-P301Ltau (S262E-OptoTau-ΔN; B) were expressed in Neuro2a cells by Lipofectamine 2000^TM^ transfection reagent. The cells were irradiated with blue light without nocodazole exposure in serum-free DMEM. After 24 hrs of irradiation, tau protein in the cells was stained with SNAP-Cell TMR-Star (magenta) (left panel). Clustering tau was counted (right panel).

(C,D) Neuro2 a cells were transfected with S262E-OptoTau-ΔN and differentiated by retinoic acid. The cells were exposed blue light for 1 hr. (C) The cells were stained with SNAP-Cell TMR-Star (magenta) and immunostained with an antibody for βIII-tubulin, a differentiation marker (green). (D) Tau phosphorylation at S422 residues was detected with reducing condition SDS-PAGE western blotting.

All error bars indicate mean ± SD. *p < 0.05 and **** < 0.0001 by Welch’s t-test (A,B) or Student’s t-test (D). W/O, without. The white bar is 10 μm.

**Figure S7**

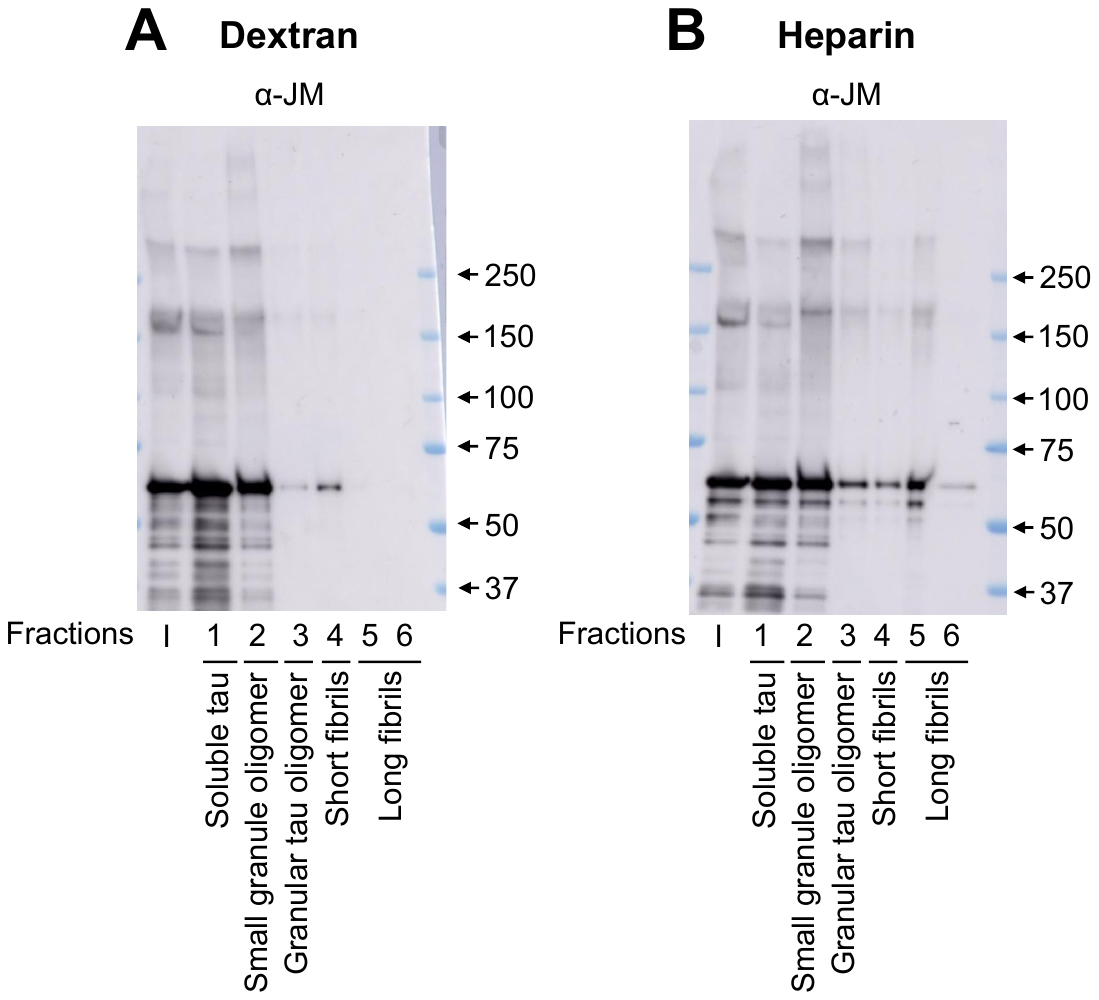

**Figure S7. Dextran-treated recombinant tau forms granular tau oligomers but not long fibrils.**

Wild-type human full-length recombinant tau (2N4R isoform) was polymerized by 0.12 mg/ml dextran (A) (Sigma Aldrich-Merck, 31404-5G-F) or 0.12 mg/ml heparin (B) in 30 mM Tris (pH=7.4) for 72 hrs at 37 ℃ according to previously described procedures ^1^ with minor modifications. After the aggregated tau mixture was separated into 6 fractions by sucrose density gradient centrifugation ^2,3^, the fractions were suspended in Laemmli sample buffer containing 3-mercapto-1,2-propanediol, and boiled. Tau protein was detected using SDS-PAGE western blotting with rabbit polyclonal pan-tau JM antibody. Tau was observed in fractions 2, 3, and 4 but not in 5 and 6 when treated with dextran (A). Heparin-treated tau was observed in all fractions (B). These indicate that dextran can form tau oligomers to short fibrils but not long fibrils, unlike heparin. I, input.

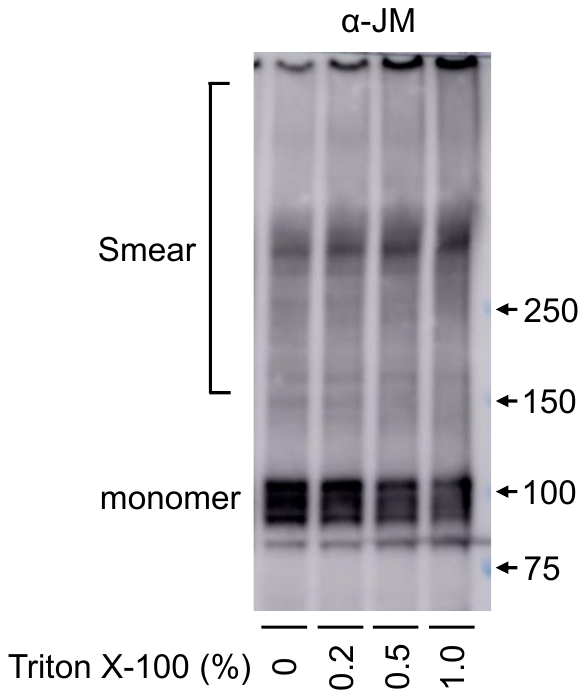

**Figure S8**

**Figure S8. Aggregates of OptoTau-ΔN are not solubilized by Triton X-100.**

Neuro2a cells expressing OptoTau-ΔN were exposed to nocodazole and blue light for 20 hrs. After the cells were homogenized in 250 mM sucrose solution and centrifuged (3000xg, 10 min, 4℃), supernatants were treated with 0.2-1.0% TritonX-100 for 20 minutes at 4 ℃ to solubilize cell organelles. However, the 1% Triton X-100 treatment did not abolish the smear band formed by blue light irradiation. Thus, the tau aggregates formed by blue light are resistant to Triton X-100.

**Figure S9**

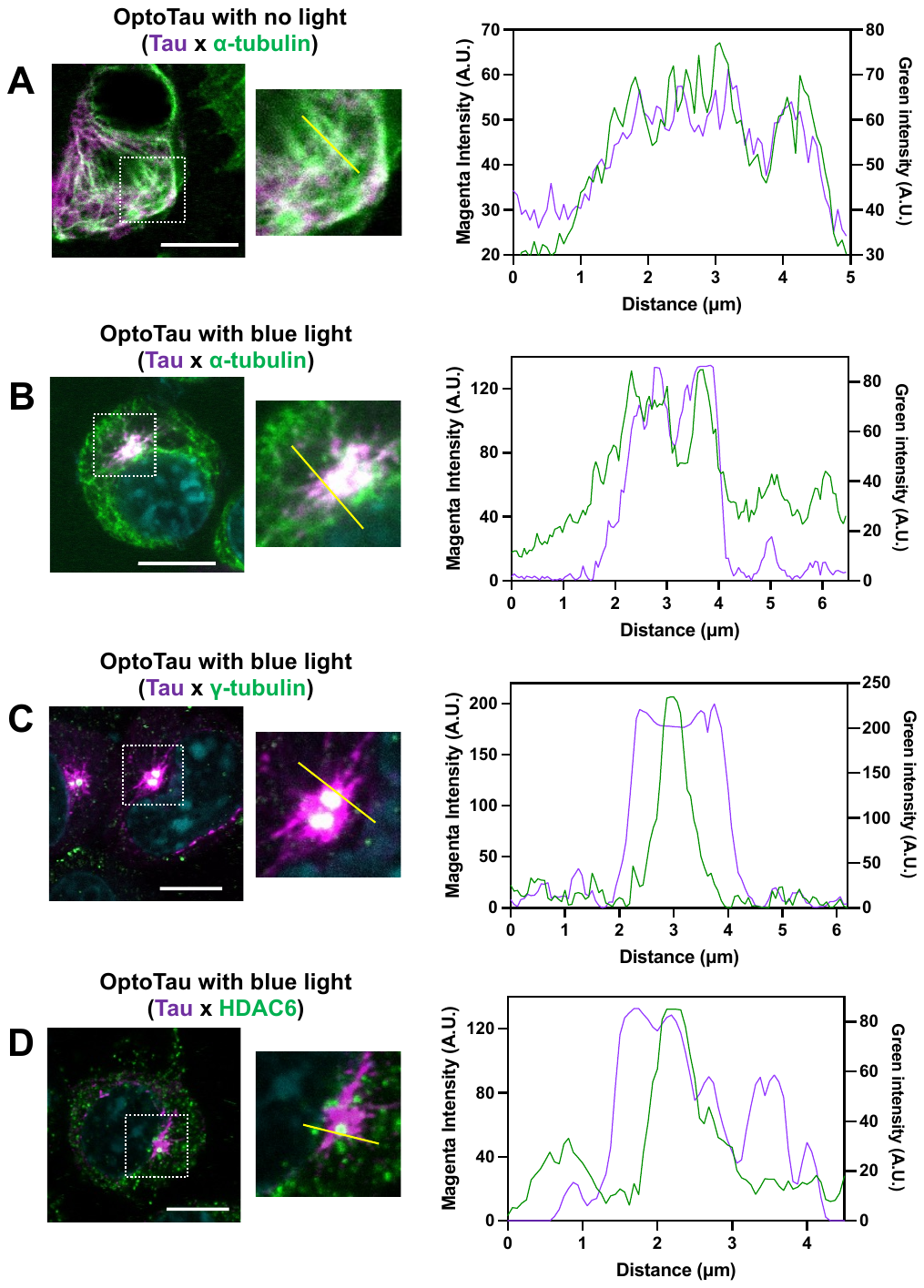

**Figure S9 continued**

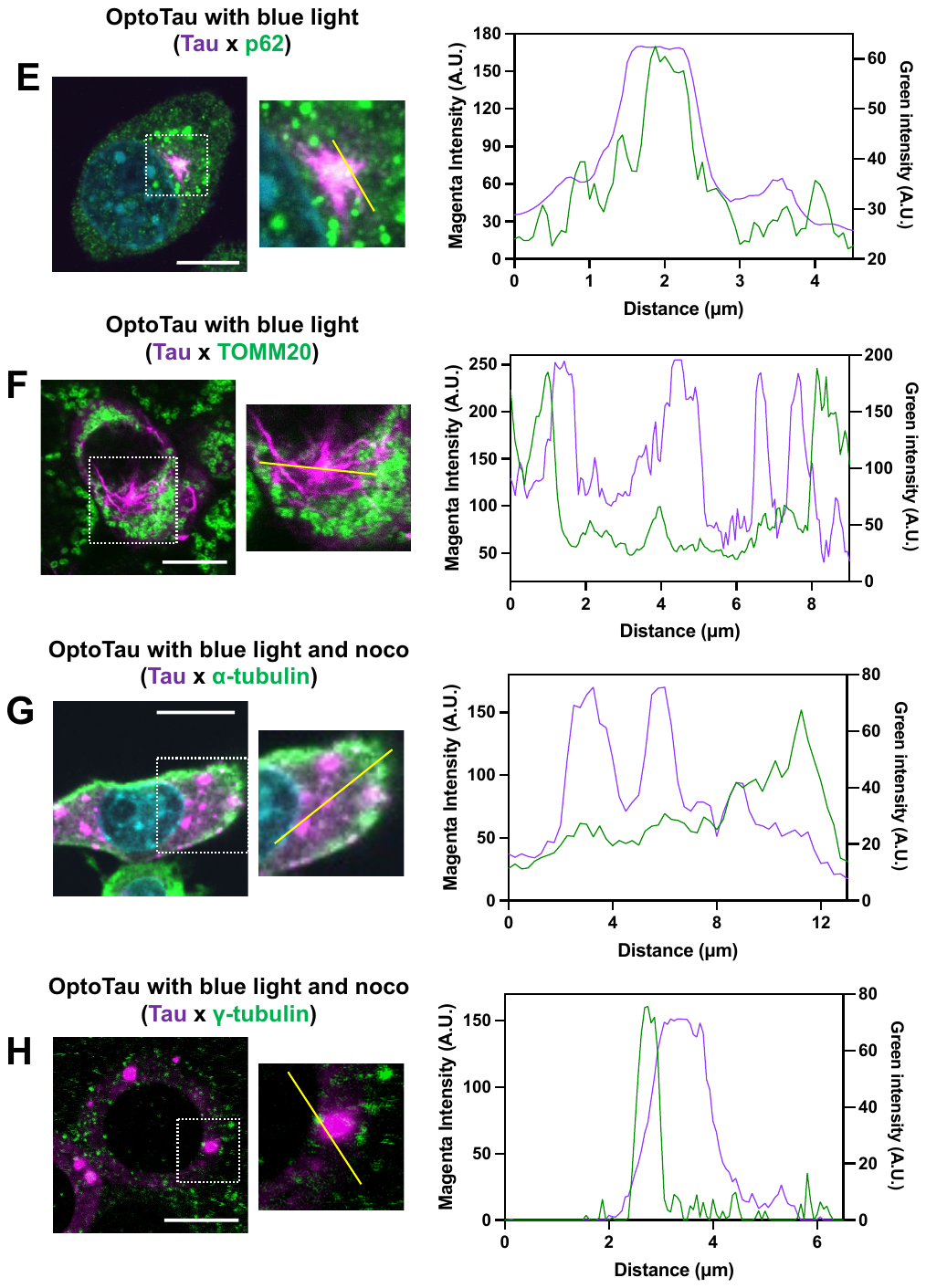

**Figure S9 continued**

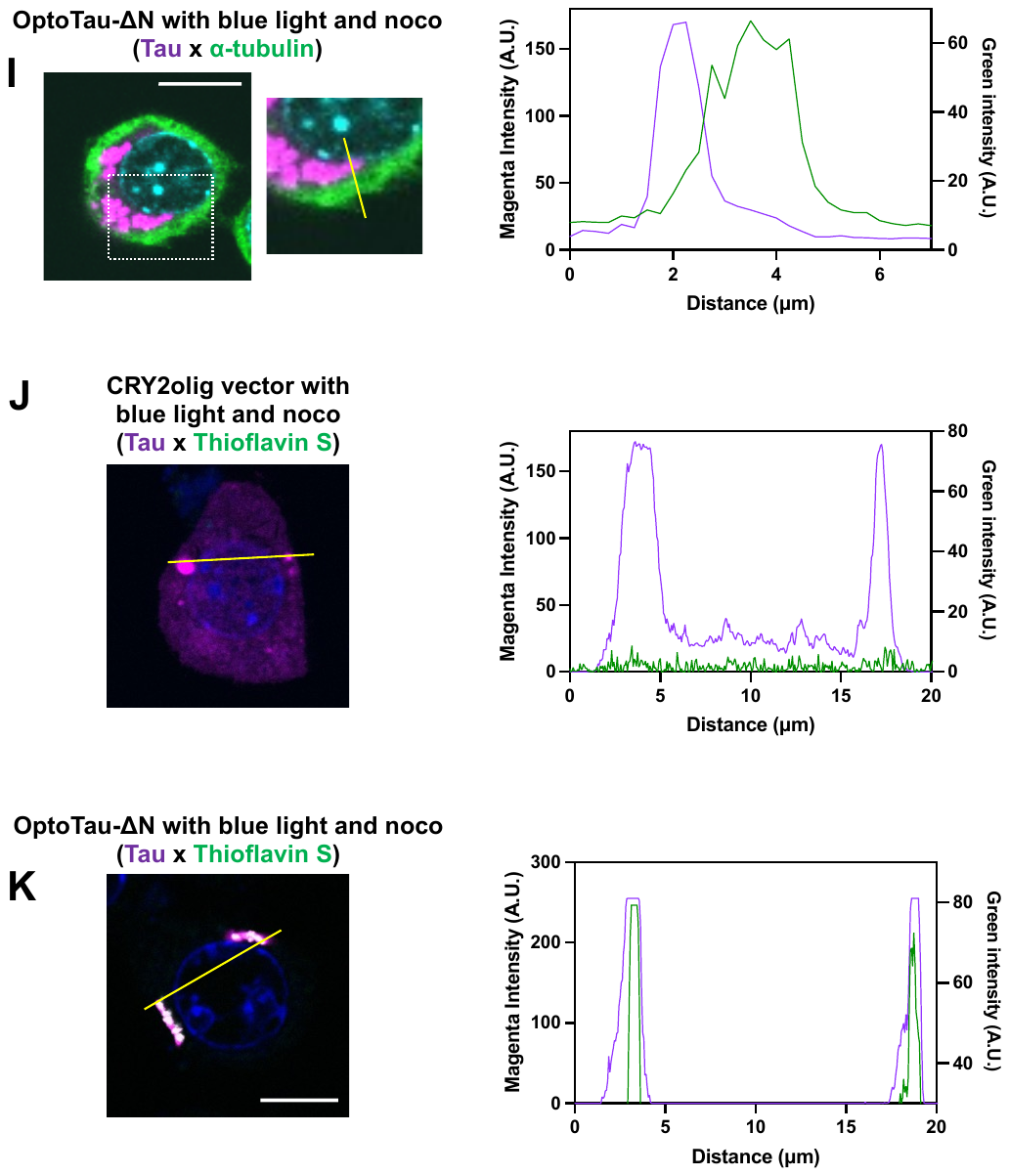

**Figure S9 continued**

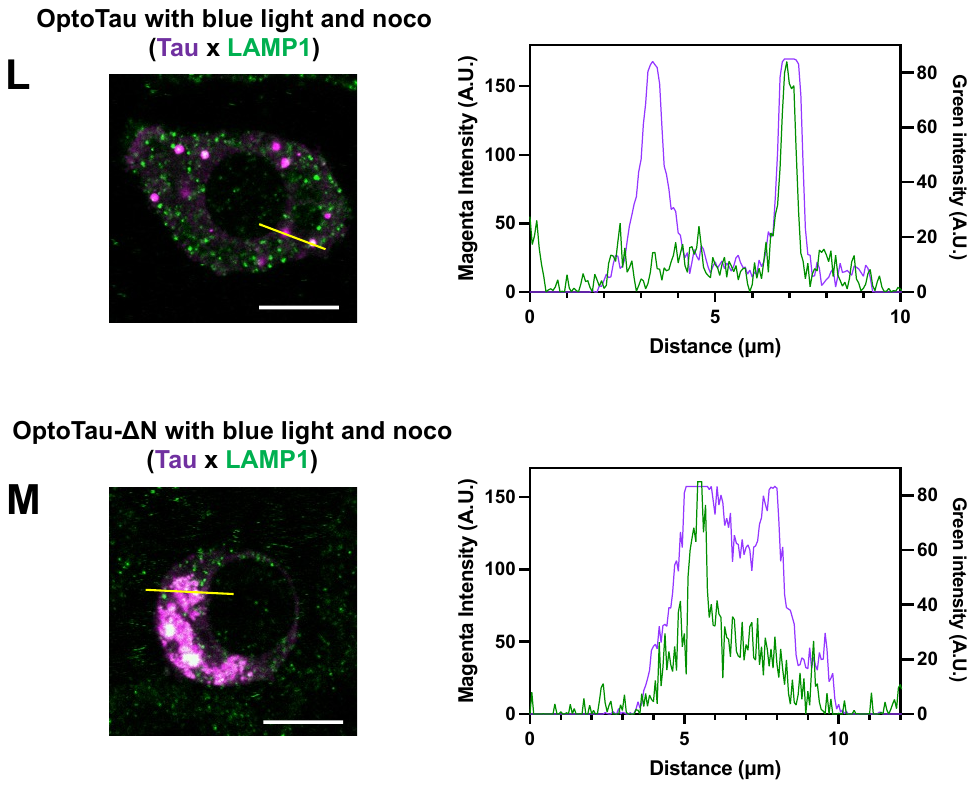

**Figure S9. Line profiles to quantify colocalizations in the images by microscopy.** Neuro2a cells were transfected with OptoTau (A-H,L) and OptoTau-ΔN (I,K,M), and were exposed to blue light (B-M) with (G-M) or without (A-F) 10 μM nocodazole. The cells were stained with SNAP-Cell TMR-Star to detect tau (magenta), and immuno-stained with α-tubulin antibody (A,B,G,I), γ-tubulin (C,H), HDAC6 (D), p62 (E), TOMM20 (F), Thioflavin S (J,K) LAMP1(L,M) (green). Merged Images were shown in left panel. The middle panels (A-I) show expanded images in the dotted white square. Intensity of the fluorescence on yellow lines were analyzed by imageJ to make line profile graphs. W/O, without; noco, nocodazole. The white bar is 10 μm.

**Figure S10**

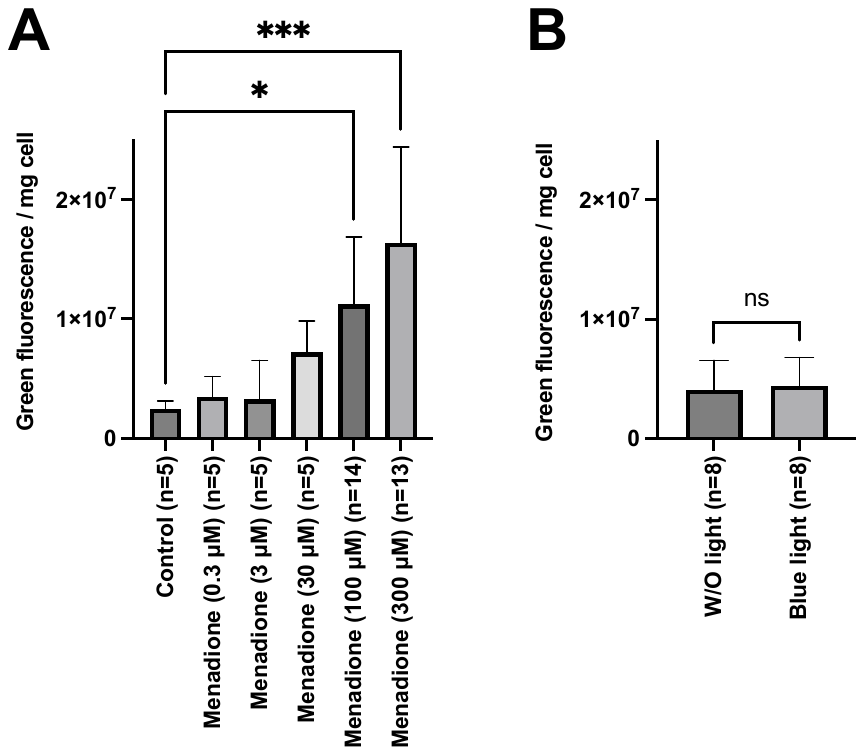

**Figure S10. The level of reactive oxygen species (ROS) under blue light exposure.** Neuro2a cells at 2.0 x 10^4^ cells were seeded on a 96-well black plate (greiner, 675096) coated with poly-D-lysine. The cells were treated with 0.3-300 μM menadione and 0.3% DMSO (control) for 1 hr (A), or exposed to blue light and W/O light for 24 hrs (B), two days after seeding. Menadione was used as a positive control to generate ROS. ROS levels were measured using CellROX Green Reagent (Thermo Fisher Scientific, C10444) with a plate reader at 460/40 nm excitation and 520/25 nm emission. After measuring ROS levels, the cells were solubilized in 1% TritonX-100 and the total protein concentration was determined using the BCA method. The intensity of the green fluorescence was normalized to the corresponding amount of total protein. All error bars indicate mean ± SD. *p < 0.05, *** < 0.001 by one-way analysis of variance and Dunnett's multiple comparisons test (A) or Student’s t-test (B). W/O, without.

**Figure S11**

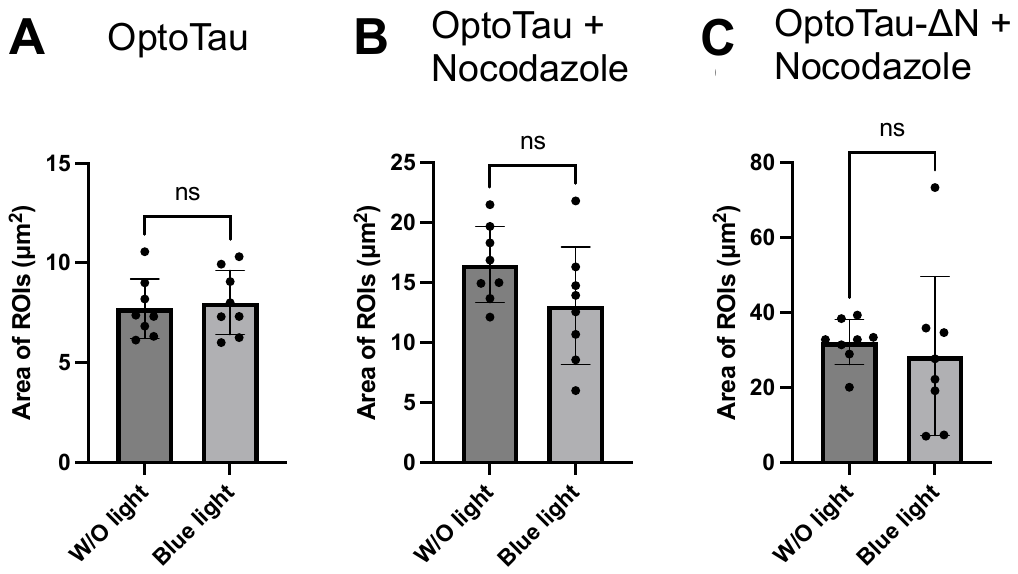

**Figure S11. No difference in area of ROIs between W/O light and blue light exposures.**

Neuro2a cells expressing OptoTau (A,B) and OptoTau-ΔN (C) were exposed with (B,C) or without (A) nocodazole. The cells were stained with SNAP-Cell TMR-Star to detect OptoTau (magenta). ROIs were set up in the images (Figs. 1D, 2E, 4D) according to the *Microscopy observation and image analysis* (see METHOD DETAILS). The intensity of tau was measured within these ROIs.

**Legends of the Supplemental Video**

**Supplemental Video 1. Imaging of OptoTau induced clustering under treatment of 1,6-hexanediol.** Neuro2a cells were transfected with OptoTau, and were exposed to blue light and 10 μM nocodazole. The cells were stained with SNAP-Cell TMR-Star, and treated with 1,6-hexanediol. The images were obtained at 0, 10, 30 and 60 minutes. Fluorescence of OptoTau induced clustering was attenuated by 1,6-hexanediol.

**Supplemental Video 2. Imaging of OptoTau-ΔN induced clustering under treatment of 1,6-hexanediol.** Neuro2a cells were transfected with OptoTau-ΔN, and were exposed to blue light and 10 μM nocodazole. The cells were stained with SNAP-Cell TMR-Star, and treated with 1,6-hexanediol. The images were obtained at 0, 10, 30 and 60 minutes. Fluorescence of OptoTau-ΔN induced clustering was not attenuated by 1,6-hexanediol.
